# Anisogamy drives diverse patterns of sex-role divergence

**DOI:** 10.64898/2026.09.26.754688

**Authors:** Xiaoyan long, Jonathan M. Henshaw

## Abstract

Male and female animals often differ remarkably in their reproductive behaviours, typified by competing males and caring females. One influential explanation traces sex-role divergence to anisogamy: sex differences in gamete size. This explanation remains controversial, because existing models have not fully resolved how anisogamy leads to secondary sex-role divergence, and exceptions to typical patterns are empirically well-documented. Here, we develop mathematical models linking anisogamy to secondary sex-role evolution, showing that anisogamy drives sex-role divergence along contrasting evolutionary trajectories depending on ecological context. When the sexes differ only in gamete size, male-biased mating competition and offspring care evolve under ‘scramble competition’, whereas uncertain paternity shifts the population towards typical patterns. By contrast, anisogamy generally drives typical sex roles under ‘contest competition’, although sex-role reversal can emerge as an alternative stable state when care must be apportioned among offspring. Anisogamy can thus explain the striking diversity of sex differences across taxa.

## Introduction

In the animal kingdom, males and females differ strikingly in their reproductive strategies. Females typically provide most or all parental care, whereas males compete intensely for mating opportunities by developing weapons, ornaments, or other traits that increase access to mates^1–4^. Understanding how these typical sex roles evolve has stimulated decades of theoretical and empirical work. One of the most influential explanations derives from Robert Trivers^3^, who argued that sex-role divergence ultimately originates from anisogamy, the gamete-size dimorphism that biologically defines the sexes, whereby females produce relatively fewer and larger eggs, while males produce smaller and more numerous sperm. According to Trivers, because females already invest substantially in gamete production, they risk losing a greater initial investment if they abandon the brood and are therefore favoured to continue investing in extended forms of parental care, i.e. post-zygotic care. This makes females the more limiting reproductive resource, intensifying male competition for access to females, which in turn reduces male investment in offspring care and thereby reinforces female-biased care.

This classical verbal explanation, in which anisogamy initiates a self-reinforcing divergence between the sexes, shaped textbook accounts and classroom lectures on sex role evolution. However, Trivers’ argument has also faced sharp criticism. First, critics have argued that it commits the “sunk cost” (or “Concorde”) fallacy, because optimal decisions should depend on future costs and benefits rather than past investment *per se*^5^. Second, the causal relationships between parental care and sexual selection are not trivial, owing to complex interactions between parental and mating strategies^6,7^. Third, this explanation does not readily account for exceptions to typical sex roles, such as species in which males provide the majority of parental care (as in many amphibians and teleost fishes^8–10^), or females compete intensely for mating opportunities^11,12^. Although Trivers also discussed species exhibiting complete sex-role reversal, characterised by caring males and competing females, he explained these cases by arguing that sufficiently high male post-zygotic care can outweigh the initial female gametic investment, thereby favouring stronger female competition for access to males. However, this explanation does not account for why males evolve greater post-zygotic care in the first place. Thus, it remains largely unclear why sex roles become reversed in some taxa but not others when the only initial sex difference is gamete type.

Following these critiques, considerable theoretical effort has been devoted to exploring whether and how an initial asymmetry at the gamete level, i.e. anisogamy, generates subsequent asymmetries in sex roles at the organism level. A central challenge in this literature is to understand the mechanisms through which initial asymmetries in gametic investment give rise to pronounced sex differences in extended forms of parental care. Several authors have argued that anisogamy alone cannot explain sex-role divergence in parental care; instead, any link between anisogamy and female-biased care is indirect, mediated by secondary effects of anisogamy such as stronger male mating competition or lower certainty of parentage in males^13,14^. Others have shown that, even in the absence of these secondary factors, the coevolution of care effort and care ability can readily promote sex-role divergence^15^; however, the direction of divergence (female care vs male care) depends on pre-existing sex differences in the costs and benefits of care, leaving open whether anisogamy itself can determine which sex evolves a greater ability to provide more care.

Surprisingly, despite efforts to formalise the link between anisogamy and female-biased care, anisogamy itself has typically been modelled only indirectly^16^. For instance, Fromhage and Jennions^17^ used small initial differences in parental investment as a proxy for anisogamy, showing that such initial asymmetries can be amplified through positive evolutionary feedback when parental effort and sexually selected traits are allowed to coevolve. In contrast, Long and Weissing^7^ specifically incorporated sex-specific pre-mating investment and showed that the “Trivers effect” can be recovered even under random mating: the sex with greater pre-mating investment (e.g. females take longer to replenish eggs than males take to replenish sperm) typically tends to evolve higher post-zygotic parental care. Intriguingly, Iyer and colleagues^18^ showed that explicitly linking sex differences to gamete types does not inevitably favour female care: when anisogamy is represented by limiting eggs and effectively abundant sperm, sex-biased care, when it evolves at all, tends to be male-biased, particularly under synchronous mating. This overall suggests a more complex causal link between anisogamy and post-zygotic parental care than Trivers and others originally envisaged.

While the direct causal pathway from anisogamy to post-zygotic care remains unresolved, recent theoretical work has formally linked anisogamy to the evolution of sex differences in mating competition^16^, supporting Darwin’s and Bateman’s original insights^1,2^. By explicitly incorporating anisogamy into models, assuming that females produce larger and therefore fewer gametes than males, Lehtonen and colleagues showed that anisogamy alone can ultimately drive males to compete more intensely for mating opportunities^19^, and cause male reproductive success to increase more steeply with the number of mating partners (i.e. steeper ‘Bateman gradients’ in males^20^). Given the sustained controversy over the role of anisogamy^21–24^, these models provide an important foundation for extending formal theory beyond mating competition towards a broader account of sex role evolution that also incorporates parental care. This extension is particularly important because parental and mating strategies often interact in complex and non-trivial ways^6^, while the pathway linking anisogamy-related investment to post-zygotic care is itself intricate^7^. Moreover, existing theory does not explain how sex-role reversal can evolve when the sexes only differ in gamete size. Consequently, the overall role of anisogamy in explaining broader secondary sex differences remains poorly understood, despite its central place in sex-role theory^16^. This uncertainty has fuelled broader critiques of the anisogamy-based view of sex roles, with some researchers arguing that sex-role theory as a whole is fundamentally flawed and proposing instead that sex roles are shaped primarily by stochastic processes or external factors, such as ecological and social conditions^21,22,25^.

### Model outline

Here, we reconcile these contrasting perspectives by developing a modelling framework that formally links anisogamy to two major sex-role components: mating competition and parental care. We define sex in the model through gamete-size dimorphism directly rather than categorising individuals as “male” or “female” arbitrarily or via non-gametic traits, as is common in previous models. Thus, females produce eggs of fixed size *m*_*f*_, whereas males produce sperm of fixed size *m*_*m*_, with eggs assumed to be always larger than sperm (*m*_*f*_ > *m*_*m*_). Because our aim is to examine whether and how anisogamy shapes sex role evolution, no other asymmetries between the sexes are assumed unless stated otherwise. To model the coevolution of reproductive strategies, we assume that each individual is endowed at birth with a fixed resource budget, *R*, allocated across three activities: gamete production, mating competition, and post-zygotic parental care (Fig. 1a). Resource allocation is determined by the heritable traits *x*_*f*_ and *y*_*f*_ in females and *x*_*m*_ and *y*_*m*_ in males. A female with these trait values dedicates a proportion *c*_*f*_ = *x*_*f*_ of her resources to mating competition, *p*_*f*_ = *y*_*f*_(1 − *x*_*f*_) to parental care and the remainder *g*_*f*_ = 1 − *c*_*f*_ − *p*_*f*_ to gamete production. Male allocation is analogous. All four heritable traits (*x*_*f*_, *y*_*f*_, *x*_*m*_, *y*_*m*_) are carried by individuals of both sexes, but only expressed in the appropriate sex.

**Fig. 1:**
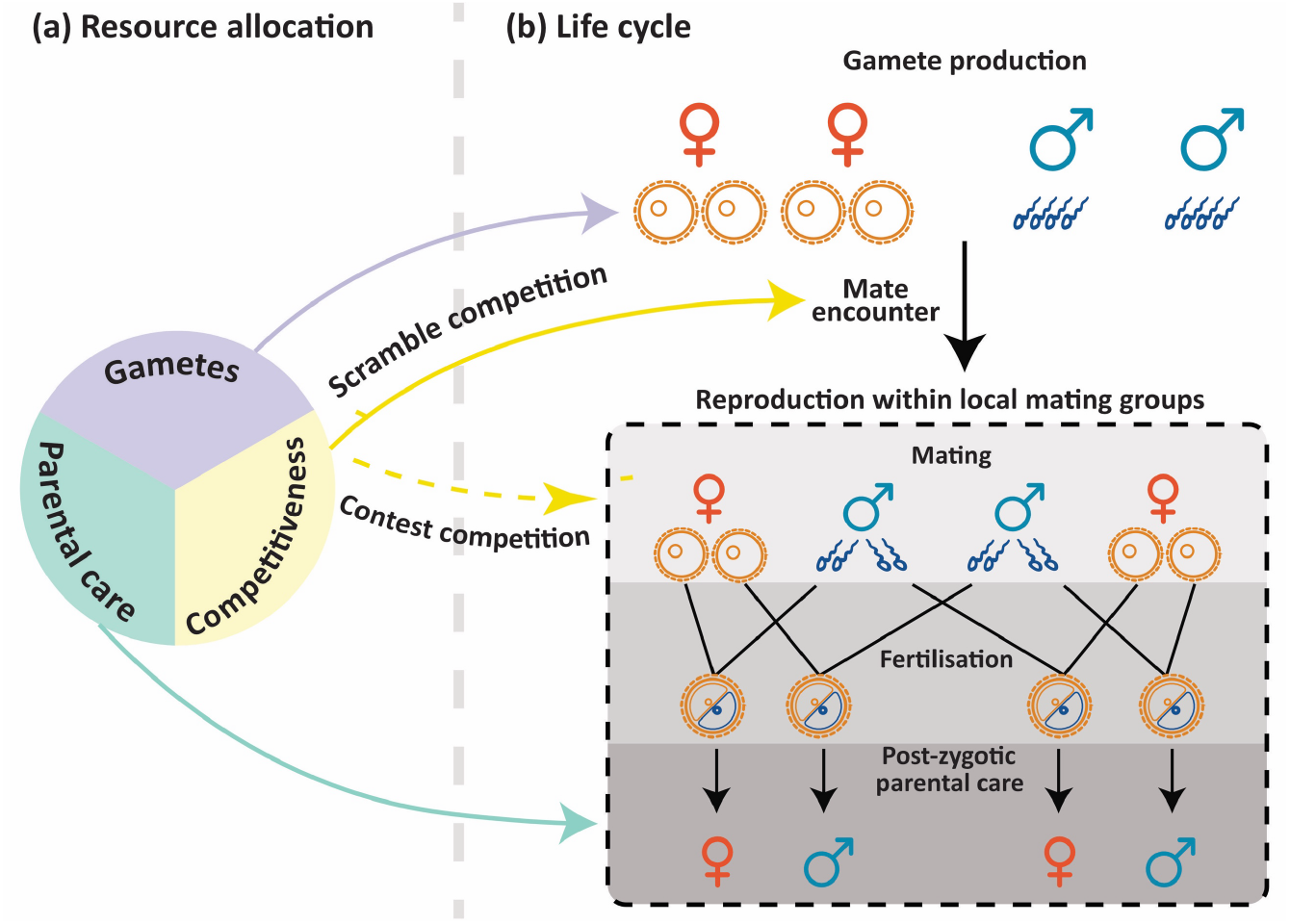
Model schematic. **(a)** Each individual possesses a fixed amount of resources that are allocated across gamete production (eggs for females, sperm for males), mating competition, and post-zygotic parental care. Sex-specific resource allocation strategies are evolving traits. **(b)** The life cycle is as follows: First, individuals produce gametes according to their allocation strategy. Females produce larger (and therefore typically fewer) gametes than males to reflect anisogamy. Greater individual allocation to gamete production yields higher gamete numbers. After gamete production, individuals enter local mating groups to seek mating opportunities. Under the ‘scramble competition’ scenario, individuals allocate resources to traits that increase mate encounter rates (e.g. rapid movement or call/pheromone production), such that higher competitiveness improves their ability to access more local groups and thereby encounter more mates; mating within each group occurs randomly. Under the ‘contest competition’ scenario, individuals enter a fixed number of groups, but mating is structured by direct competitive interactions within those groups: individuals that invest more in competitive traits (e.g. weapons or ornaments) win more contests over the opposite sex and thus gain a greater share of mating success. Fertilisation is then determined by the numbers of eggs and sperm present, with fertilisation efficiency assumed to be high. Fertilised eggs develop into zygotes, after which parental investment in post-zygotic care determines offspring survival. An individual’s fitness is proportional to its total number of surviving offspring.

Based on resource allocation strategies, individuals proceed through the life cycle depicted in Figure 1b. Each individual allocates a proportion of its resources to gamete production, with greater investment yielding more gametes. As a result, each female produces *n*_egg_ = *Rg*_*f*_/*m*_*f*_ eggs and each male produces *n*_sperm_ = *Rg*_*m*_/*m*_*m*_ sperm.

After gamete production, individuals enter local mating groups to search for mating opportunities. Here, we consider two mating competition scenarios to capture the different mechanisms through which individuals can compete for access to mating partners or to the resources those partners provide^26^. One common form is ‘scramble competition’, which is widespread among arthropods^27,28^, anurans^29^ and mammals^30^. In scramble competition, individuals compete through traits and behaviours that increase their mate encounter rates, such as active searching for mates via rapid movement or long-distance signalling through calls or pheromones; thus, mating success depends largely on how efficiently individuals locate potential partners or make themselves detectable to them^27,31,32^. In contrast, sexual selection also involves direct ‘contest competition’ in many animals, whereby individuals develop weapons such as horns or antlers used to fight rivals, or ornaments that attract mating partners^26,32^. In these cases, successful competitors can monopolise access to multiple mates; mating success therefore depends mainly on gaining priority access to mates through direct contests or mate attraction. Accordingly, under the ‘scramble competition’ scenario, greater investment in competitiveness is assumed to allow individuals to enter more local groups and thereby encounter more potential mating opportunities. Under the ‘contest competition’ scenario, by contrast, individuals invest in traits used directly in competitive interactions within local groups, enabling more competitive individuals to monopolise access to a greater number of partners (see below for details of both scenarios). Note that we use the terms ‘scramble competition’ and ‘contest competition’ more broadly than is typical in the literature. The most important difference between these two scenarios in our model is that mating success is driven by relative competitiveness under contest competition, whereas absolute competitiveness is key under scramble competition.

Within a local group, the total number of fertilisations is determined by the total numbers of the two gamete types that are effectively available for fertilisation, with fertilisation efficiency assumed to be high unless stated otherwise (see Supplementary Information and Supplementary Fig. 1 for details). Once an egg and a sperm fuse to form a zygote, an offspring is produced, with sex determined randomly with an equal probability of being male or female. After producing offspring, parents provide post-zygotic care, and offspring survival depends on the total amount of care received from both parents.

Building on these general assumptions, we use adaptive dynamics to analyse the evolution of resource allocation across the three reproductive activities in males and females under different mating competition scenarios, assuming that the primary sex difference lies in gamete size (see Methods for more details on the invasion fitness analysis). We first consider a baseline model in which (i) parents care only for their own offspring, thereby ensuring complete paternity certainty, and (ii) parental care is non-depreciable^4,33^: the same amount of care can be provided to multiple offspring simultaneously (e.g. nest oxygenation or defense^33^), so per-offspring care benefits do not decline with the number of siblings. We then extend the model to consider paternity uncertainty, as commonly occurs in internally fertilising species^34^, and depreciable parental care^4^, in which care allocated to one offspring is no longer available to benefit another (e.g. food provisioning). Overall, our model provides mathematical evidence that gamete-size dimorphism can indeed explain the widespread tendency for males to compete more strongly than females for fertilisations, while females provide most post-zygotic care. Intriguingly, depending on the ecological context, anisogamy can also explain exceptions to these typical sex roles, including cases of complete sex-role reversal, where females compete intensely for mating opportunities and males provide most offspring care.

## Results

### Anisogamy drives male-biased mating competition and offspring care under scramble competition

To represent scramble competition in our model, we assume that greater investment allows individuals to visit more local mating groups and thus encounter more potential partners (Fig. 1). Specifically, the average number of local groups a resident individual can enter is given by *θ*_*f*_ = *kc*_*f*_*R* for females and *θ*_*m*_ = *kc*_*m*_*R* for males, where *k* is a scaling parameter that determines how easily individuals can locate and enter local mating groups. Each local mating group then contains *n*_*f*_ = (1 − *r*)*θ*_*f*_*d* females and *n*_*m*_ = *rθ*_*m*_*d* males. Here, *r* denotes the adult sex ratio (ASR, the proportion of males in the adult population), with a default value of 0.5 given that no sex-specific life-history differences are assumed in the current model. The scaling parameter *d* determines the expected group size when each individual enter a single group, thereby capturing environmental or ecological constraints on group formation. Note that when *θ*_*f*_ ≠ *θ*_*m*_, the numbers of males and females within each group can differ, such that the local operational sex ratio (OSR, the proportion of males among those individuals in the local group that are available to mate) is not necessarily 0.5. Fertilisation within each local group is random, with an individual’s fertilisation success depending on the number of gametes they produce, but not on their competitiveness (see Methods and Supplementary Information for details). Key parameters and variables are summarised in Table 1.

**Table 1.** Overview of key notation.

| Symbol | Meaning |
| --- | --- |
| $R$ | Resource budget for each individual |
| $x_f, y_f, x_m, y_m$ | Evolving resource allocation strategies |
| $c_f, c_m$ | The fraction of resources allocated to mating competition; $c_f = x_f, c_m = x_m$ |
| $p_f, p_m$ | The fraction of resources allocated to post-zygotic parental care;<br>$p_f = y_f(1 - x_f), p_m = y_m(1 - x_m)$ |
| $g_f, g_m$ | The fraction of resources allocated to gamete production;<br>$g_f = 1 - c_f - p_f, g_m = 1 - c_m - p_m$ |
| $m_f, m_m$ | Gamete size (for eggs, sperm) |
| $k$ | Mate encounter efficiency (scramble competition scenario) |
| $d$ | Group size coefficient, representing the expected size of local groups when each individual is present in only one group (scramble competition scenario) |
| $r$ | Adult sex ratio (ASR) |
| $\delta$ | “Winner-takes-all” effect (contest competition scenario) |

The subscripts $f$ and $m$ denote females and males, respectively.

Figure 2a shows the evolutionary trajectories of female and male resource allocation strategies across the three reproductive activities under scramble competition, at an intermediate expected group size (*d* = 10). The two sexes differ inherently only in gamete size: *m*_*f*_ = 0.1, *m*_*m*_ = 0.01. Our adaptive-dynamics model predicts that, regardless of initial conditions, the two sexes evolve distinct allocation strategies. Males allocate most of their resources to scramble competition (i.e. mate searching or long-distance signalling traits; Fig. 2a_2_), followed by post-zygotic parental care (Fig. 2a_3_) and sperm production (Fig. 2a_1_). In contrast, females allocate the majority of their resources to egg production (Fig. 2a_1_), followed by scramble competition (Fig. 2a_2_), and the least to parental care (Fig. 2a_3_). Overall, males engage in slightly higher levels of scramble competition and contribute a greater share of post-zygotic care than females, whereas females invest more heavily in gamete production. The magnitude of sex differences in resource allocation across the three reproductive strategies increases with the degree of gamete size dimorphism; however, this effect eventually levels off, such that further increases in gamete-size dimorphism produce little additional divergence between the sexes (Fig. 2b, also see Fig. 4b). This is consistent with recent theoretical predictions that anisogamy generates a broad sex difference in sexual selection, but that, once anisogamy is already pronounced, further variation in the degree of gamete-size dimorphism is not expected to closely track variation in sexual-selection intensity^35^. This perspective helps reconcile two apparently contrasting empirical patterns: the degree of anisogamy show little correlation with variation in sexual selection intensity across animal taxa^36^, yet males nevertheless tend to compete more intensely for mating opportunities than females^12^. Our results extend this pattern beyond sexual selection alone, showing that a similar saturation effect can also apply to broader sex-role divergence involving gamete production and post-zygotic parental care.

**Fig. 2:**
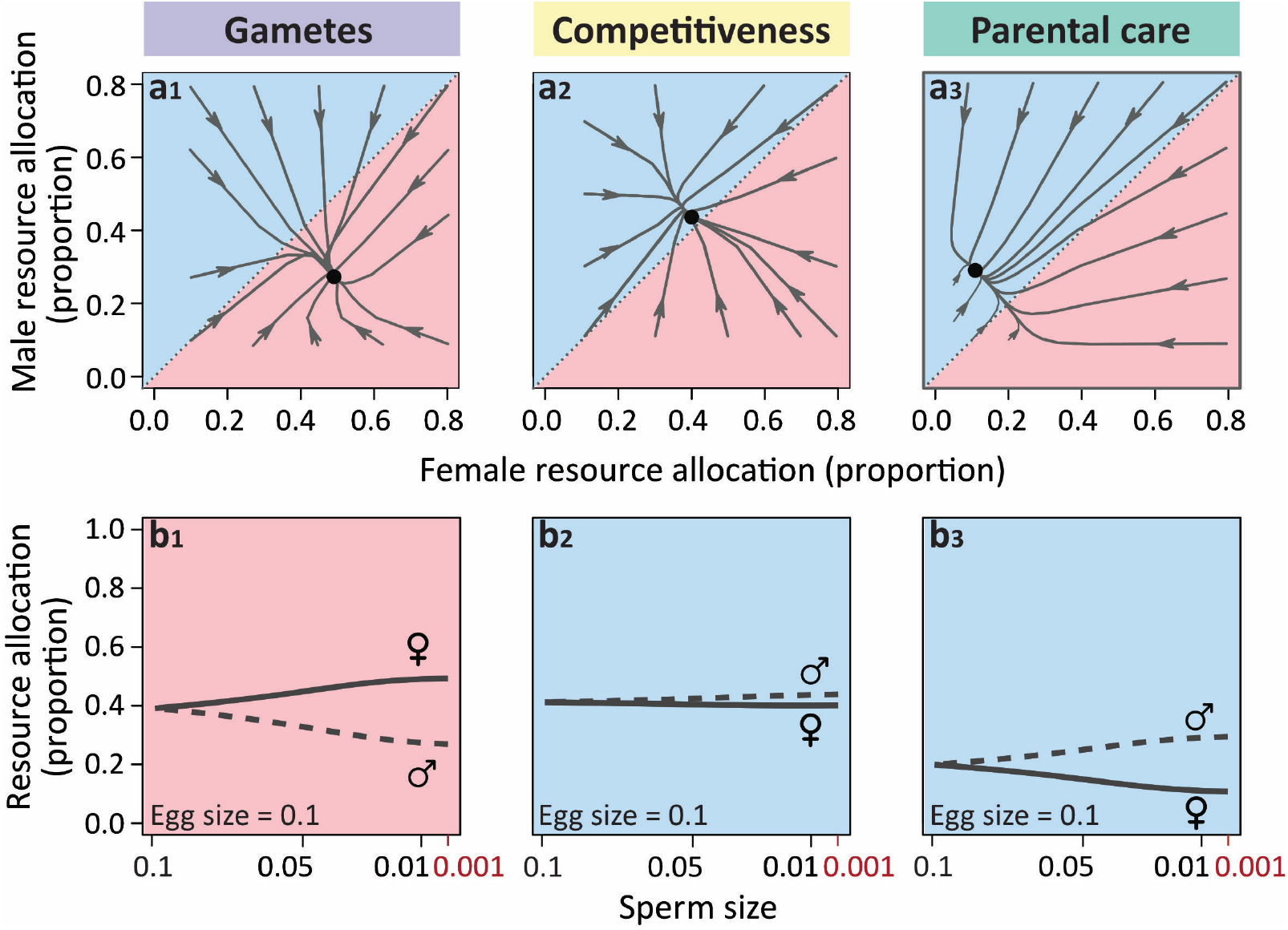
Resource allocation strategies under scramble competition. The panels show how anisogamy influences the proportion of resources allocated to gamete production, mating competition, and post-zygotic parental care under scramble competition. **(a)** Evolutionary trajectories of female and male allocation strategies starting with different initial conditions, with egg size *m*_*f*_ = 0.1 and sperm size *m*_*m*_ = 0.01. Arrows indicate the direction of evolution from various starting values of the evolving resource-allocation traits, and black dots denote equilibrium points. At equilibrium, females allocate more resources to gamete production (a_1_), whereas males allocate more to competitive traits for scramble competition (a_2_) and post-zygotic care (a_3_). **(b)** Sex differences in resource allocation, as shown in (a), increase with the degree of anisogamy (i.e., the ratio of sperm to egg size). Solid dark grey lines represent equilibrium female allocation strategies under different parameter values, whereas dashed dark grey lines represent equilibrium male allocation strategies. Throughout the manuscript, light red shading indicates female-biased allocation, and light blue shading indicates male-biased allocation. All populations were initialised at *x*_*f*_ = *y*_*f*_ = *x*_*m*_ = *y*_*m*_ = 0.1 in (b). Other key parameter values for both panels: *R* = 1, *k* = 1, *d* = 10, *r* = 0.5.

Our evolutionary outcomes are strikingly different from what has often been predicted, particularly because previous theory did not explicitly incorporate gamete production or the mechanisms of mating competition. The evolutionary logic behind this pattern can be understood as follows: Female gametes are substantially larger than male gametes, with the result that any fixed investment yields far fewer eggs than sperm. When fertilisation is highly efficient, as assumed here, this ensures that most eggs are fertilised. As a result, the fitness value of eggs is roughly proportional to the number produced, all else being equal. Therefore, selection favours females that allocate most of their resources to egg production, while maintaining a certain level of investment in competitiveness to reduce the risk of failing to enter any local mating group, as individuals that enter no group encounter no mates and achieve zero reproductive success. In contrast, most sperm are not fertilised. When local groups are small, this leads to diminishing fitness returns on sperm production due to local sperm competition (i.e., wasteful competition among an individual male’s own sperm)^37^. This incentivises males to redirect investment away from gamete production and towards other traits. In particular, the low level of female care in this scenario creates a selection environment favouring greater male investment in post-zygotic care, without which offspring survival would be low. Males consequently invest relatively heavily in both scramble competition and parental care.

This interpretation is further supported by model predictions when varying additional ecological factors. While keeping all other assumptions identical to the setup used in Figure 2a, we explored several scenarios in which one ecological parameter was varied at a time. First, we show that as fertilisation efficiency decreases, sex differences in resource allocation across the three reproductive traits become less pronounced (Supplementary Fig. 2). This occurs because declining fertilisation efficiency reduces saturation in the relationship between sperm number and fertilisation success, thereby linearising fitness returns on sperm production^19,38,39^. As a result, males redirect resources away from mating competition and parental care and towards sperm production. Females are then favoured to increase investment in post-zygotic care to compensate for reduced male care, reallocating resources away from egg production and mating competition.

Second, varying the expected group size parameter *d* produces a similar pattern (Supplementary Fig. 3). As *d* increases, more individuals aggregate within each local mating group, reducing the need for mate-encounter effort in both sexes (Supplementary Fig. 3a_2_), while intensifying sperm competition (Supplementary Fig. 3a_1_), since more males compete for the limited eggs. This again makes fitness returns on sperm production more nearly linear, shifting male investment towards sperm production and away from the other two traits. Females are consequently favoured to increase parental care at the expense of egg production, and the resulting decline in egg availability further intensifies sperm competition within local groups. As a result, secondary sex differences decrease as expected local group size increases.

Third, when offspring can survive in the absence of care (see Methods for details), increasing baseline survival reduces offspring dependence on parental care, thereby lowering parental investment by both sexes (Supplementary Fig. 4). In response, female care decreases to zero first, leading to a transition from male-biased to male-only care, and ultimately to no parental care when baseline survival is sufficiently high (Supplementary Fig. 4c). This may help explain why uniparental care is more common in nature than our baseline model might suggest, given that offspring in many taxa may retain some chance of survival even without parental care^40–42^. When parental care is no longer essential for offspring survival, both sexes reallocate the freed resources to gamete production and mating competition. Consistent with Figure 2a, females prioritise egg production while maintaining just enough investment in competitiveness to ensure encounters with males. Males, by contrast, allocate more resources to scramble competition than to sperm competition, given that the realised local group size is small and local sperm competition is consequently strong (Supplementary Fig. 4a,b).

### Anisogamy drives typical sex roles under scramble competition in the presence of uncertain paternity

In contrast to the assumptions of the baseline model, uncertain paternity is common in many animals, particularly in those with internal fertilisation^34,43,44^. In such cases, only females can typically be certain of their parentage, and genetic analyses often reveal that some offspring are sired by extra-pair males, even in socially monogamous species^43,44^. Sex differences in parentage certainty are widely thought to explain, at least in part, why males typically provide less post-zygotic parental care than females, with numerous theoretical studies showing that male care declines as paternity becomes less certain^13,14,17^.

To investigate how reproductive strategies under scramble competition evolve with paternity uncertainty, we introduce an additional source of sex asymmetry beyond anisogamy. Specifically, we assume that a fraction *η* of a brood is sired by the focal male, whereas the remaining 1 − *η* is sired by other males, reflecting uncertain paternity. In a population with male care, a focal male’s fitness therefore comprises two components: (i) offspring he sires and subsequently cares for, and (ii) offspring he sires but that are cared for by other males. Under these assumptions, uncertain paternity does not affect the fitness of resident males, but it does influence the invasion fitness of a rare mutant male through both components (see Methods for details).

Under scramble competition, male care declines with increasingly uncertain paternity and ultimately reaches zero, while female care increases correspondingly (Fig. 3c). This shift generates a transition from male-biased care when paternity is certain to female-biased and eventually female-only care as paternal uncertainty increases. As male care decreases (Fig. 3c), males reallocate resources toward both sperm production and scramble competition (Fig. 3a,b). In contrast, females respond by increasing their investment in post-zygotic care (Fig. 3c), accompanied by reduced allocation to egg production and mating competition traits (Fig. 3a,b). Consequently, females produce fewer eggs and participate in fewer local mating groups, becoming more a limiting resource for reproduction. This, in turn, intensifies sexual selection on males, reinforcing the emergence of typical sex roles characterised by intense male mating competition and female-biased (or even female-only) post-zygotic care.

**Fig. 3:**
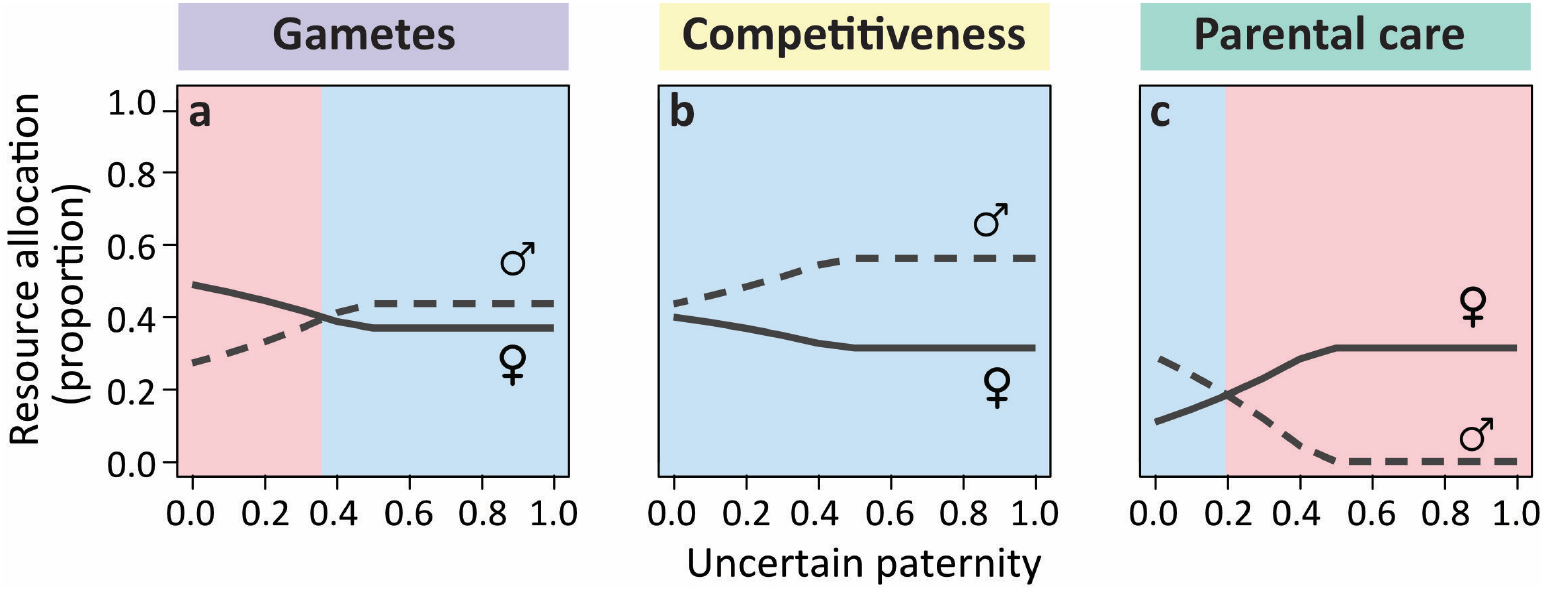
Resource allocation under scramble competition with uncertain paternity. Females and males allocate resources to **(a)** gamete production, **(b)** scramble competition, and **(c)** post-zygotic parental care in an internally fertilising population with potentially uncertain paternity. A fraction *η* of each brood is sired by the focal male, whereas the remaining fraction 1 − *η* is assumed to be sired by other males. All populations were initialised at *x*_*f*_ = *y*_*f*_ = *x*_*m*_ = *y*_*m*_ = 0.1. Other key parameter values: *R* = 1, *m*_*f*_ = 0.1, *m*_*m*_ = 0.01, *k* = 1, *d* = 10, *r* = 0.5.

### Anisogamy drives typical sex roles under contest competition

In the contest competition scenario, we assume that individuals enter a fixed number of local mating groups, which therefore have a fixed, unbiased sex ratio in terms of both the ASR and OSR. In this scenario, however, mating is no longer random within each group: individuals with higher competitiveness (e.g. larger weapons or more elaborate ornaments) gain a larger share of mating opportunities by excluding rivals. Consider a mutant male with competitive trait ĉ_*m*_ who competes for access to females within a local group that otherwise consists of resident individuals. The effective number of rival males with whom it competes for fertilisations is assumed to be 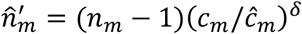, where *n* is the number of males in the group, *c*_*m*_ is the resource allocation to competitiveness among resident males within the local group, and *δ* represents a “winner-takes-all” effect that determines the strength of sexual selection. Higher values of *δ* generate a stronger skew in mating success, with mating opportunities increasingly concentrated among the most competitive individuals. The effective number of rivals for a mutant female 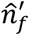 is defined analogously (see Methods for further details on the invasion fitness analysis).

Compared to the scramble competition scenario, this form of mating competition leads to fundamentally different patterns of resource allocation when the only difference between the sexes is gamete size (Fig. 4). Starting from various initial allocation conditions, the model consistently converges to typical sex roles (Fig. 4a), characterised by intense male-male contest competition (Fig. 4a_2_) and strongly female-biased post-zygotic care (Fig. 4a_3_). Because both sexes are assumed to enter a fixed number of local mating groups (here with each individual entering one group on average), females can reliably encounter males and achieve fertilisation without investing in competition (Fig. 4a_2_). By contrast, males produce large numbers of sperm, and their reproductive success increases with the number of eggs they fertilise. Thus, males increase their share of mating success through substantial investment in contest competition (Fig. 4a_2_), with the remaining resources allocated to sperm production (Fig. 4a_1_). In response, females provide nearly all post-zygotic parental care (Fig. 4a_3_), while their residual resource goes into egg production (Fig. 4a_1_). As expected, sex differences in resource allocation strategies increase markedly with the degree of anisogamy, but eventually reach a plateau once anisogamy becomes sufficiently pronounced (Fig. 4b).

**Fig. 4:**
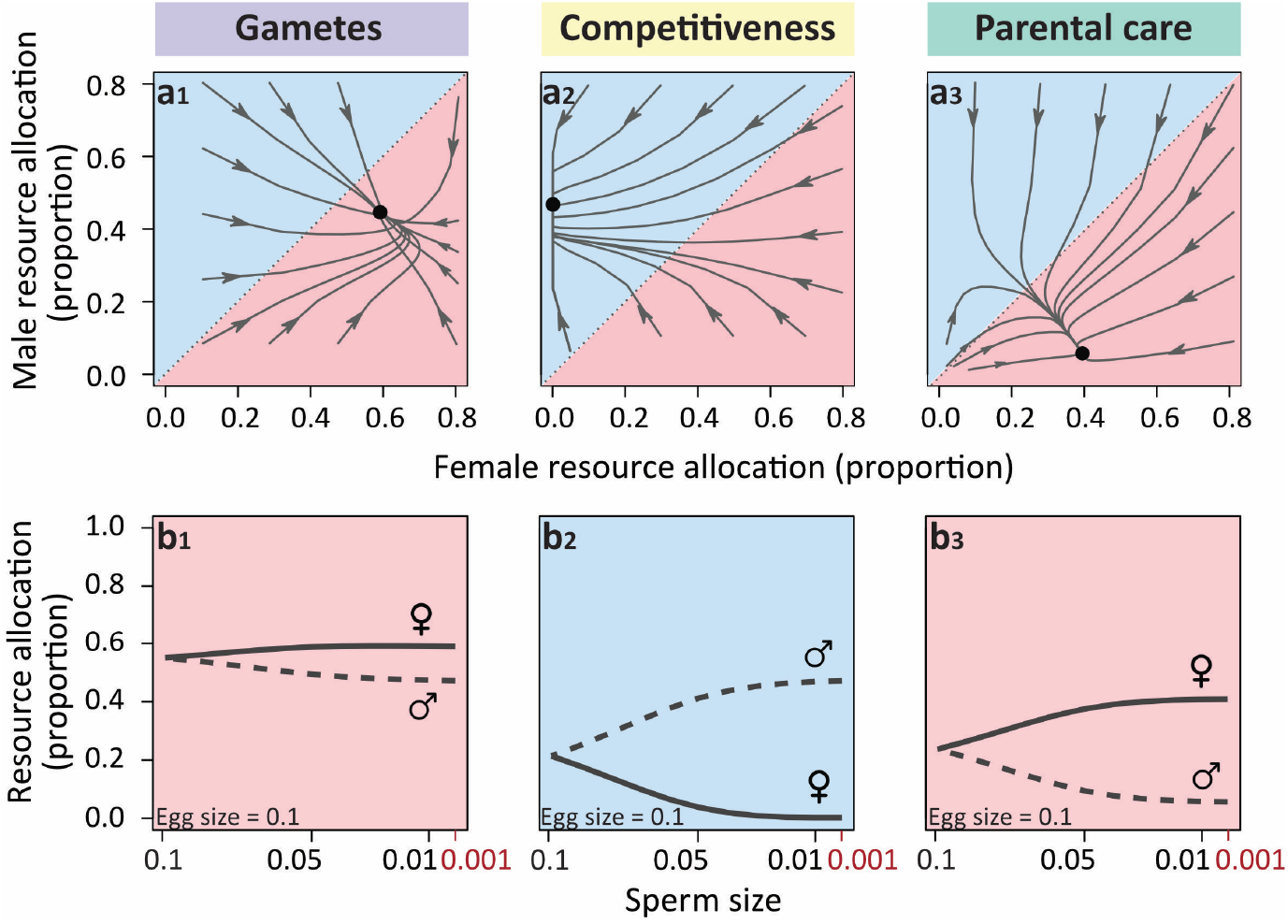
Resource allocation strategies under contest competition. The panels depict **(a)** the evolutionary trajectories of female and male resource allocation strategies under contest competition with anisogamy (*m*_*f*_ = 0.1, *m*_*m*_ = 0.01), and **(b)** how sex differences in resource allocation change with the degree of anisogamy (i.e., the egg–sperm size ratio). Because scramble competition is absent, here each individual is assumed to enter one local mating group on average. There are thus five males and five females in each local group. In (a), arrows indicate the direction of evolution from various starting values, and black dots denote equilibrium points. In (b), all populations were initialised at *x*_*f*_ = *y*_*f*_ = *x*_*m*_ = *y*_*m*_ = 0.1. Other parameter values: *R* = 1, *d* = 10, *r* = 0.5, *δ* = 1; since the number of groups each individual enters is not affected by competitiveness, *θ*_*f*_ = *θ*_*m*_ = 1.

Interestingly, local group size has an important influence on resource allocation strategies under contest competition (Supplementary Fig. 5). The results shown in Fig. 4 are derived from moderately large group sizes (five males and five females per group). When local group size is small, the model recovers patterns similar to those observed under scramble competition, with males providing most parental care and sometimes investing slightly more than females in mating competition. For example, when each local group contains only one male and one female, there is no need for investment in contest competition or additional sperm production driven by local sperm competition. As local group size increases, however, competition among males intensifies, favouring greater investment in competitive traits to increase mating success. Consequently, the system transitions towards typical sex roles as shown in Fig. 4a. The effect of paternity uncertainty on sex role evolution here is similar to that under scramble competition: males further reduce their investment in parental care, even though care levels are already very low when paternity is certain (since the evolutionary outcomes are qualitatively very similar to those shown in Fig. 4a, these results are not shown).

### ‘Winner-takes-all’ effect exaggerates sex-role divergence under contest competition

In some animal species, contest competition can be extremely strong, leading to high reproductive skew in the more competitive sex. For example, in lekking species, individuals of one sex (typically males) compete intensely for access to mates, and only a small number of successful individuals monopolise the majority of mating opportunities^45^. Such “winner-takes-all” patterns are well documented in certain bird and mammal species^46–48^.

As the winner-takes-all parameter in our model increases, males invest more heavily in traits associated with contest competition (Fig. 5b). Because fewer males monopolise access to females within each group, sperm competition is reduced, leading to decreased investment in sperm production (Fig. 5a). Females, in turn, are selected to provide all post-zygotic parental care (Fig. 5c), and allocate their remaining resources to egg production (Fig. 5a). When the ‘winner-takes-all’ effect is weak (here *δ* < 0.5), the fitness returns from contest competition are limited. Under these conditions, decreasing *δ* leads males to invest less in competitive traits and to reallocate some of the freed resources to post-zygotic care (Fig. 5b,c). In response, females reduce their investment in parental care, thereby freeing additional resources for egg production (Fig. 5a).

**Fig. 5:**
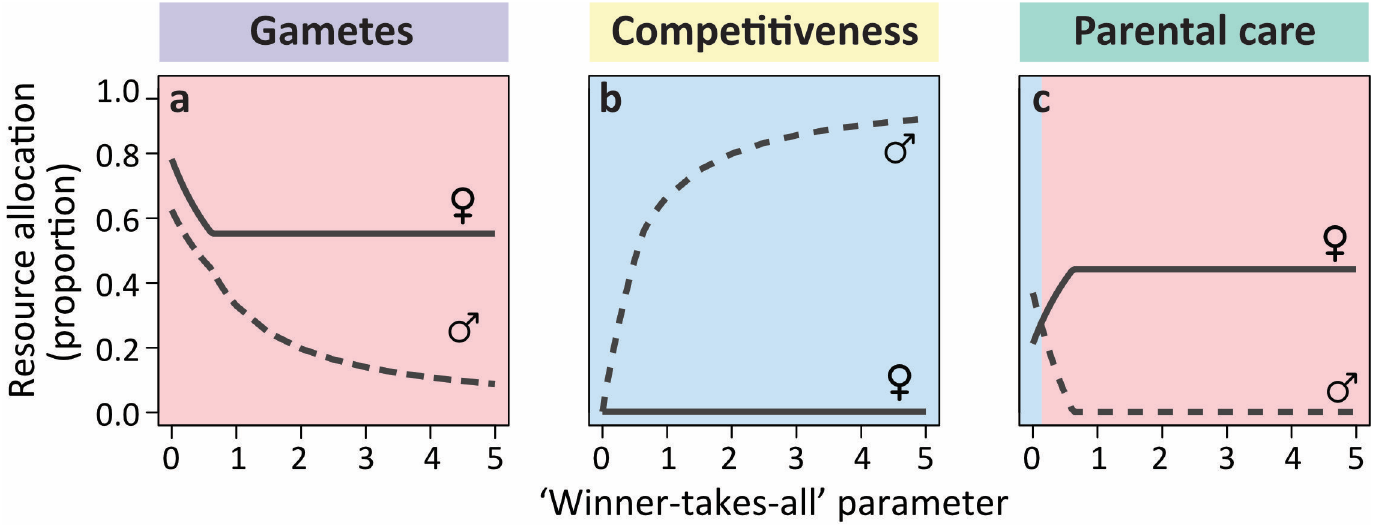
‘Winner-takes-all’ competition exaggerates sex-role divergence. The figure shows how increasing the strength of intrasexual competition (i.e., the winner-takes-all effect, *δ*) influences resource allocation to **(a)** gamete production, **(b)** contest competition, and **(c)** post-zygotic parental care. As intrasexual competition becomes stronger, successful males monopolise a greater number of mating partners. This increases the fitness returns to investment in competitive traits, leading males to allocate even more resources to mating competition while reducing investment in sperm production and parental care. As males withdraw from parental care, females compensate by providing all post-zygotic care. Because sperm remains sufficient for fertilisation within each group, females do not engage in competition for mates and therefore allocate no resources to mating competition, instead investing the remaining resources in egg production. All populations were initialised at *x*_*f*_ = *y*_*f*_ = *x*_*m*_ = *y*_*m*_ = 0.1. Other parameter values: *R* = 1, *m*_*f*_ = 0.1, *m*_*m*_ = 0.01, *d* = 10, *r* = 0.5, *θ*_*f*_ = *θ*_*m*_ = 1.

### The evolution of alternative sex roles under contest competition when parental care is depreciable

Up to this point, we have assumed non-depreciable parental care, whereby a parent can care for multiple offspring simultaneously without reducing the benefit received by each offspring^4,33^. This form of care is common in groups such as teleost fishes^33^, where parental activities (e.g. nest oxygenation or defense) are roughly equally effective regardless of brood size. In contrast, many other species, including birds and mammals, exhibit depreciable care^4^, in which parental investment (e.g. provisioning young with food) must be divided among offspring, thereby reducing the benefit per offspring as brood size increases. To incorporate depreciable post-zygotic care, we assume that each parent’s investment in care is evenly distributed among all offspring, such that the maternal and paternal care received per offspring decreases as brood size increases (see Methods for details). All other model assumptions follow those in Fig. 4.

Interestingly, when post-zygotic care is depreciable, we observe two alternative evolutionary outcomes (Fig. 6). In most cases, the model converges to typical sex roles (black points), as in the non-depreciable scenario: strong male–male competition and female-only post-zygotic parental care (Fig 6b,c). Under this equilibrium, females allocate less resources to egg production compared to the case of non-depreciable care (Fig. 6a), resulting in smaller clutch sizes. This reduction arises because producing additional offspring reduces the amount of care each offspring receives, thereby lowering their probability of survival. By contrast, when females initially exhibit greater competitiveness than males (yellow trajectories in Fig. 6) or males initially provide substantially more care than females (green trajectories), the model converges to an alternative equilibrium (red points). In this case, sex roles are completely reversed: females engage in intense contest competition, while males provide all post-zygotic parental care. Here, females invest more in egg production and thus produce larger clutches than in the conventional equilibrium (black points). This arises because females compete for access to caring males. As a result, competitive females can monopolise multiple males that invest heavily in parental care, allowing each offspring to receive sufficient care despite its depreciable nature. Notably, although these alternative resource allocation strategies emerge from different initial conditions, anisogamy continues to constrain the evolutionary outcomes. Rather than fully symmetric dynamics given different initial conditions, anisogamy biases the system toward the traditional sex role equilibrium, influencing both the direction and stability of sex-role evolution (also see Supplementary Fig. 6 for evolutionary outcomes across a broader range of initial values for the resource allocation variables *x*_*f*_, *y*_*f*_, *x*_*m*_, *y*_*m*_).

**Fig. 6:**
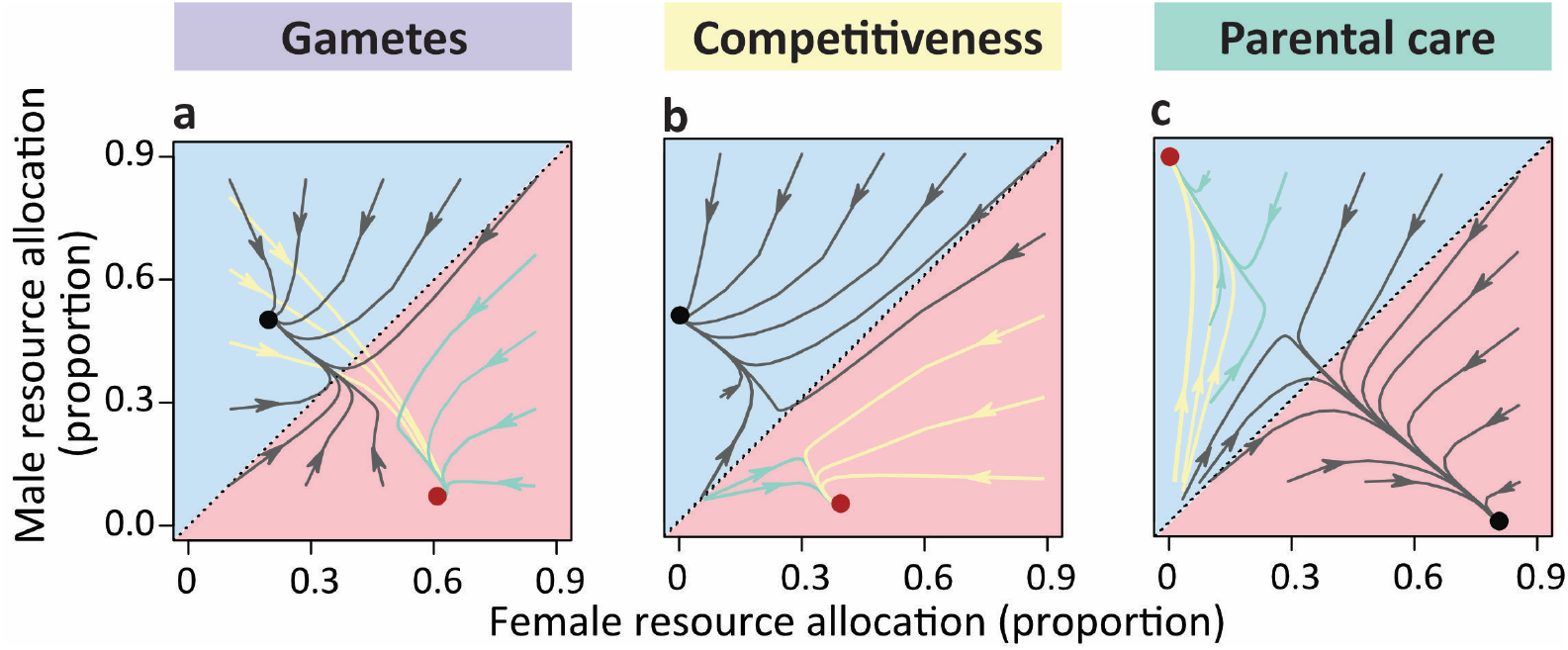
Resource allocation under contest competition with depreciable post-zygotic care. The graphs depict the evolutionary trajectories of female and male resource allocation to **(a)** gamete production, **(b)** contest competition and **(c)** post-zygotic care, under anisogamy (*m*_*f*_ = 0.1, *m*_*m*_ = 0.01) and various initial conditions. In most cases, the population converges to ‘conventional sex roles’ (black dots), characterised by strong male–male competition and female-only care. Here, males invest more in sperm production than females invest in eggs. However, when females initially engage in strong mating competition (yellow trajectories) or males initially invest heavily in parental care (green trajectories), the population instead converges to an alternative equilibrium with reversed sex roles (red dots), characterised by female–female competition and male-only care. At this equilibrium, females allocate more resources in gamete production than males. Arrows indicate the direction of evolution. Other parameter values: *R* = 1, *d* = 10, *r* = 0.5, *δ* = 1, *θ*_*f*_ = *θ*_*m*_ = 1.

We also examined the scramble competition scenario with depreciable care (Supplementary Figs. 7-9), which yielded patterns qualitatively similar to those under non-depreciable care (cf. Figs. 2 and 3; Supplementary Fig. 3), but with several important differences. First, clutch size is generally smaller, because females invest less in egg production. Second, when males provide care, females invest more in competitiveness, allowing offspring to be distributed across multiple caring males. Therefore, when parental care is depreciable, both sexes engage in mating competition, as under non-depreciable care, but for sex-specific reasons here: males compete for fertilisation opportunities over a limited number of eggs, while females compete to mate with multiple caring males.

## Discussion

Here, we present a comprehensive modelling framework that formalises the link between anisogamy and two of the most striking sex roles: mating competition and post-zygotic parental care. By explicitly defining the sexes based on gamete type and allowing for the coevolution of male and female investment in gamete production, mating competition, and parental care, we provide theoretical evidence that anisogamy can drive sex-role divergence. Intriguingly, this divergence follows contrasting evolutionary trajectories depending on ecological context. First, we show that when the sexes differ only in gamete size and post-zygotic care is non-depreciable, anisogamy can lead to different patterns of sex role evolution depending on the mechanism of mating competition. Under scramble competition, males evolve both higher competitiveness and greater investment in post-zygotic care than females (Fig. 2). By contrast, under contest competition, typical sex roles evolve, corresponding to strongly male-biased mating competition and strongly female-biased post-zygotic care (Figs. 4, 5). Second, when paternity uncertainty is introduced as an additional sex asymmetry, our model shows that males reduce their investment in parental care as uncertainty increases, consistent with previous models^13,14,17^. Under scramble competition, this reduction in male care can shift the population towards typical sex roles, with males competing more to increase mate encounter rates and females providing most or all post-zygotic care (Fig. 3). Third, when gamete-size dimorphism is the only initial asymmetry, but parental care is depreciable, the population can evolve towards either typical sex roles (with competing males and caring females), or complete sex-role reversal (with competing females and caring males) under contest competition; however, due to anisogamy, the typical sex-role equilibrium has a larger basin of attraction, so it is reached from more initial conditions than the sex-role reversal equilibrium (Fig. 6 and Supplementary Fig. 6). Note that despite the diversity of secondary sex-role characteristics, females generally allocate more resources to gamete production than males throughout our model analyses. This is perhaps unsurprising, given empirical evidence that, in most motile organisms, females produce a greater total gamete volume than males, whereas males produce more gametes overall^49^.

Our findings reconcile contrasting perspectives on the roles of anisogamy and external environments in explaining sex role evolution: anisogamy provides the initial sex asymmetry, while environmental conditions shape how this asymmetry unfolds, giving rise to diverse patterns of sex-role divergence through different evolutionary pathways. Notably, environmental conditions themselves need not necessarily generate sex-specific selection, as some critiques have suggested^21,22,24^; rather, they can shape the reproductive contexts that link gamete-size dimorphism to secondary sex-role differences. For instance, the way individuals compete for mating opportunities is suggested to depend, at least in part, on ecological conditions that determine the potential to monopolise mates or mating resources^50^.

When critical resources such as food, breeding sites, or oviposition sites are widely dispersed, potential mates are also likely to be spatially dispersed. Under these conditions, defending resource patches may be difficult or prohibitively costly, increasing the importance of scramble mating competition^51^. By contrast, when mating opportunities are spatially clumped and defensible, individuals can monopolise access to multiple mates, favouring contest competition over mates^50^. This is also supported by our model predictions when expected local group size varies, a parameter that can capture the extent to which individuals are spatially aggregated under local ecological conditions. Lower expected local mating group size reflects more dispersed mating opportunities, whereas larger local group size reflects greater local aggregation. Under scramble competition, lower expected local group size therefore favours increased allocation to mate-acquisition traits in both sexes, with a stronger effect in males due to anisogamy (Supplementary Fig. 3b). Under contest competition, however, the same parameter has a different implication: small local groups provide little opportunity to monopolise multiple mates, so contest competition remains weak or even absent; as local groups become larger, opportunities for mate monopolisation increase, thereby intensifying contest competition particularly among males whose reproductive success is limited by access to eggs (Supplementary Fig. 5). Moreover, paternity uncertainty is generally not regarded as an evolving trait in itself, but instead arises from behavioural interactions between males and females, which can be largely mediated by ecological factors^52^. Abundant food, limited breeding sites, or high population density may promote breeding aggregations, increasing opportunities for female multiple mating and sperm competition and thereby generating paternity uncertainty^43,53^. That said, the mechanisms generating paternity uncertainty can be far more complex, often resulting in substantial variation at both the population and individual levels^54^. This complexity is not captured by the current model, which therefore does not explain the diverse relationships between paternity uncertainty and male care observed in empirical systems^52^.

Given the well-established causal link between anisogamy and sexual selection^16,19,20^, and the more nuanced picture showed throughout this manuscript, an important question remains: what is the direct causal link between anisogamy and post-zygotic care when no other sex role component is considered? When males and females enter a fixed number of local mating groups, mating occurs randomly within each group, and thus, individuals do not involve in any forms of mating competition, we discover that anisogamy alone drives greater male investment in post-zygotic care, alongside greater female allocation to egg production (Supplementary Fig. 10). This resulting sex difference is especially pronounced when gamete-size dimorphism is strong (Supplementary Fig. 10a), or when local group size is small and sperm competition is therefore weak or even absent (Supplementary Fig. 10b). This contrasts sharply with Trivers’ classical view, but is consistent with previous theoretical work potentially linking anisogamy to male-biased parental care^18^. Using a recently developed feedback-analysis framework for coevolving traits^55^, we decompose evolutionary change into components that quantify how changes in one trait alter selection on another, thereby providing a more explicit account of the causal pathway from anisogamy to male-biased post-zygotic care (see Supplementary Material and Supplementary Figs. 11-13 for details): Initial sex asymmetry in gamete size does not favour greater female contribution to post-zygotic care through a self-reinforcing process; instead, it leads males to provide more post-zygotic care via dynamic interactions among traits within and between the sexes. When males and females initially allocate resources equally between gamete production and post-zygotic care but differ in gamete size, the relatively low number of eggs makes each egg especially valuable under high fertilisation efficiency. Selection therefore favours females that produce more eggs, shifting resources away from post-zygotic care. However, increasing allocation to egg production progressively weakens selection for further reductions in female care. This inhibiting feedback, arising from the trade-off between offspring number and offspring survival, would be the dominant constraint initially on further reductions in parental investment if only one sex were considered. In a two-sex system, however, the continued decline in female care also becomes a major force shaping the selection environment experienced by males. Declining female care first weakens selection for reduced male care and increased sperm production, until males begin to increase post-zygotic care and reduce sperm production; thereafter, it reinforces selection in this new direction. This suggests that sex differences in post-zygotic care can arise through a cross-trait reinforcing process, in which the evolution of male care strategies is mediated by female allocation strategies when the sexes differ only in gamete size. This quantitative decomposition of the causal pathway also provides strong interpretative support for our more complex model predictions, such as those from the baseline scramble competition scenario, where males evolve both greater investment in post-zygotic parental care and stronger mating competition (e.g. Fig. 2).

Our findings may provide plausible mechanisms for the origin and subsequent evolutionary transitions of sex roles in some animal taxa. Male-biased or male-only care has been suggested as a possible ancestral state following the transition from no care to parental care in several clades, including anurans^9,56^, teleost fishes^57,58^, and birds^59,60^. Relatively few studies have attempted to address why this transition is prevalent in these taxa. For example, male territoriality or low costs of parental care have been proposed as possible explanations^58,59^. Our study suggests, however, that these transitions need not be driven by external changes, but may instead trace back to anisogamy in lineages whose ancestors experienced little or no mating competition, or mainly scramble competition. It would therefore be insightful to reconstruct ancestral states not only for parental care, but also for mating strategies in these lineages. Beyond explaining the origins of male care, our model also sheds light on how an ancestral state of male care may shift towards diverse sex-role patterns. First, in line with findings of phylogenetic comparative studies^34,58^, transitions from male care to female care can be facilitated by changes in fertilisation mode, in that males are generally expected to have higher paternity certainty under external fertilisation, as in many fishes^34^, but lower certainty under internal fertilisation, as in many birds and mammals^43,44^. Second, when mating competition creates opportunities for successful individuals to monopolise access to partners (e.g. via weapons and/or ornaments), anisogamy can shift the population towards greater female care and stronger male competition. Given the widespread occurrence of such mating competition forms across animals^26,32^, this may help explain why typical sex roles are so common. Third, the origin of male care may also provide favourable initial conditions for complete sex-role reversal, which occurs in our model only under contest competition when parental care cannot be shared among siblings, as is often the case for complex forms of care such as food provisioning. However, if such forms of care are later lost (e.g. feeding behaviour can be lost rapidly in amphibians^10^), sex-role reversed systems can transition back towards typical sex roles. By contrast, the reverse transition, from typical sex roles to complete sex-role reversal, was not possible under the conditions examined in our model. This asymmetry may help explain why complete sex-role reversal is relatively rare in nature. Lastly, transitions from male care or female care to equal biparental care occur only under very restricted ecological conditions in our model (e.g. at the points where female and male care allocation intersect in Figs. 3, 5 and Supplementary Figs. 5, 9), consistent with the general rarity of biparental care in many animal groups^9^. Intriguingly, if male care was the ancestral state in birds, it remains puzzling why and how females in later lineages evolved to make nearly equal contributions to offspring care, given the widespread occurrence of biparental cooperation in this clade^4,9^. Although previous studies suggest that biparental care can be facilitated when parental contributions from both parents have synergistic effects (e.g. through division of labour)^7,17,61^, this may not fully explain the origin of biparental cooperation or the diverse patterns of cooperation observed across avian species^62^.

Together, these findings highlight our model as a foundation for disentangling how anisogamy shapes sex role evolution and point to several promising avenues for future work. First, although our model captures scramble and contest competition conceptually, these two forms of mating competition lead to contrasting pathways of sex role evolution. Future work could incorporate more explicit mechanisms of sexual selection, such as mate choice and coevolving sexually selected traits, considering “direct benefits”, “sexy sons”, and “good genes” variants^63,64^. This would make it possible to examine whether and how anisogamy predicts which sex evolves greater choosiness, and how this, in turn, interacts with other components of sex-role evolution. Second, species may adopt alternative reproductive strategies under different ecological conditions^65–67^, for example relying more on scramble competition when monopolising mates is difficult, but more on contest competition when mates can be guarded. By explicitly incorporating ecological conditions and allowing organisms to make decisions based on their own state and environmental context, we may clarify how exactly ecological factors mediate the link between anisogamy and sex-role evolution, and potentially explain more of the population-level and individual-level variation observed in nature. Third, a promising and exciting extension would be to allow anisogamy itself to evolve. Rather than assuming any initial difference between the sexes, as in the current work, a model starting from isogamy could explore how gamete-size dimorphism, and thus the sex itself, emerge in the first place, and whether feedbacks between evolving gamete size and subsequent sex roles ultimately help explain the origin and diversification of sex differences^16^.

## Methods

In our adaptive-dynamics model, invasion fitness is defined as the expected number of offspring produced by a rare mutant male or female that survive to reproductive maturity in a genetically monomorphic resident population. To derive invasion fitness for each scenario considered in our study, we first calculate the number of fertilisations attributable to each resident individual and the survival probability of each resulting zygote. We then introduce a rare mutant to evaluate its expected reproductive output in that resident background. Supplementary Table 1 provides an overview of all symbols used in the analysis.

### Fertilisation within each local mating group in the resident population

In each local mating group, the number of fertilisations depends on the eggs and sperm effectively available for fertilisation. As shown above, each female produces *n*_egg_ = *Rg*_*f*_/*m*_*f*_ eggs and each male produces *n*_sperm_ = *Rg*_*m*_/*m*_*m*_ sperm, where *R* is the resource budget, *g*_*f*_ and *g*_*m*_ are the proportions of resources allocated to egg and sperm production, respectively, and *m*_*f*_ and *m*_*m*_ are egg and sperm size, respectively. An individual’s gametes are then distributed equally among the local mating groups it enters. Under contest competition, individuals are assumed to enter a fixed number of local groups, with the average number of groups entered set to one by default for both females and males (*θ*_*f*_ = *θ*_*m*_ = 1). Under scramble competition, however, the number of groups entered depends on competitiveness: more competitive individuals enter more groups and thereby encounter more mating opportunities. The average number of groups entered is then given by *θ*_*f*_ = *kc*_*f*_*R* for females and *θ*_*m*_ = *kc*_*m*_*R* for males, where *k* scales how easily individuals locate and enter local mating groups, and *c*_*f*_ and *c*_*m*_ denote the proportions of resources allocated to mating competition in females and males, respectively.

In principle, we would like to consider variation around these averages, in particular to account for the special case where an individual enters zero mating groups^68^. Rather than considering this complexity explicitly, we simplify by assuming that each individual enters either zero or a fixed positive number of groups, and that the composition of groups entered is always the same. Supposing that the number of groups that a female enters follows a Poisson distribution with mean *θ*_*f*_, the probability that she enters zero mating groups is 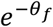. The mean number of mating groups that she enters, given that she enters at least one, is then 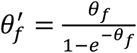. Assuming that she enters exactly 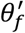 groups, she contributes 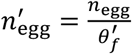eggs to each group. Males similarly contribute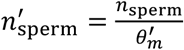to each group they participate in, where 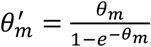. Consequently, the total numbers of eggs and sperm in each local group are 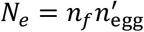 and 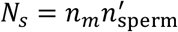, respectively. Here, *n*_*f*_ = (1 − *r*)*θ*_*f*_ *d* and *n*_*m*_ = *rθ*_*m*_*d* denote the numbers of females and males within each group, where *r* indicates the adult sex ratio, and *d* scales the expected local group size.

Given the total numbers of sperm and eggs within each local mating group, the total fertilization is assumed to be 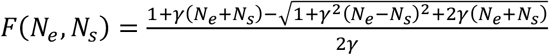 (see Supplementary Information for details of the derivation), where *N*_*e*_ and *N*_*s*_ denote egg and sperm numbers, respectively, and *γ* measures fertilisation efficiency (*γ* > 0). Throughout the manuscript, fertilisation efficiency is set to *γ* = 1 by default unless stated otherwise. This value ensures high fertilisation efficiency (see Supplementary Fig. 1). In a resident population, each female fertilises 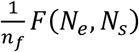eggs within a local group. Across all groups she enters, her total number of fertilised eggs is therefore 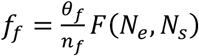.

Similarly, a resident male fertilises a total of 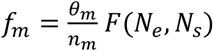 his sperm. Notably, (1 − *r*)*f*_*f*_ = *rf*_*m*_, consistent with the Fisher condition. In the resident population, the fertilisation process is similar under scramble and contest competition, except that the expected number of local groups entered is fixed under contest competition but depends on absolute competitiveness under scramble competition. When a rare mutant is introduced, however, the numbers of eggs and sperm effectively available for fertilisation within a group differ between the two scenarios, as described below in the derivation of invasion fitness.

### Offspring survival in the resident population

Once zygotes are formed through fertilisation, parents provide post-zygotic care according to their resource allocation strategies. Offspring survival depends on the total care received and is given by 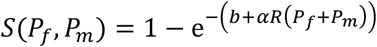, where *b* is a baseline survival parameter that determines offspring survival in the absence of parental care (note that *S*(0,0) = 1 − e^−*b*^) and *α* measures the effectiveness of parental care in enhancing offspring survival.

Throughout the manuscript, we set *b* = 0 and *α* = 1 unless otherwise stated. This means that offspring require some post-zygotic parental care to survive. Here, *P*_*f*_ and *P*_*m*_ denote the proportions of resources allocated to maternal and paternal care that are received per offspring, respectively. When care is non-depreciable, parental investment is not diluted across offspring, such that *P*_*f*_ = *p*_*f*_ and *P*_*m*_ = *p*_*m*_. In contrast, when care is depreciable, resources allocated to parental care are divided among offspring. The total actual number of offspring produced by a resident female and a resident male are given by 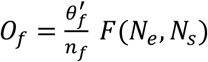 and 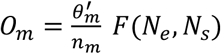, respectively. As a result, each female and male allocate a proportion *P*_*f*_ = *p*_*f*_/*O*_*f*_ and *P*_*m*_ = *p*_*m*_/*O*_*m*_ of their resources to each offspring.

Accordingly, the fitness of a resident female and male are given by *w*_*f*_ = *f*_*f*_*S*(*P*_*f*_, *P*_*m*_) and *w*_*m*_ = *f*_*m*_*S*(*P*_*f*_, *P*_*m*_), respectively.

### Invasion fitness under scramble competition

Consider a rare mutant male with traits 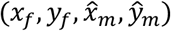 arising in a resident population (note that mutant traits are denoted by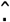 throughout). This male allocates a fraction 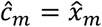 of resources to scramble competition, 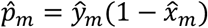 to post-zygotic care, and 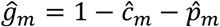 to sperm production. Thus, it produces 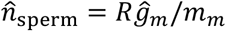 sperm and enters 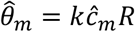 local mating groups. As described above, it enters at least one group with probability 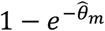 conditional on entering at least one, it accesses 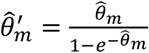 groups. Therefore, it contributes 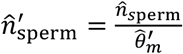 sperm to each group he enters. The total number of sperm competing for fertilisations in such a group is then 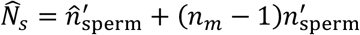, where 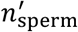 denotes the number of sperm contributed by each resident male. Consequently, total fertilisation within such a group is given by 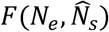, and the fraction of sperm contributed by the mutant male is 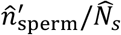.

The mutant male therefore fertilises a total of 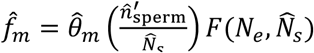 eggs across all groups he enters. The invasion fitness of the mutant male, defined as the expected number of surviving offspring, is thus 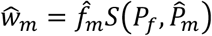, Analogously, the invasion fitness of a mutant female is given by 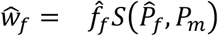, where 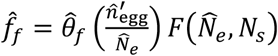, with 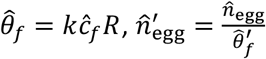, and 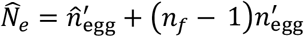.

### Invasion fitness under contest competition

Consider again a rare mutant male in a resident population, in this case under the contest competition scenario. It produces 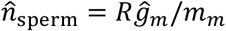 sperm and enters a fixed number of local mating groups. As described above, under contest competition, individuals that allocate more resources to mating competition can monopolise a larger share of mating opportunities. Accordingly, the effective number of rival males the mutant competes with for fertilisations is assumed to be 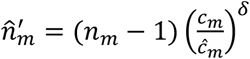. The total number of sperm that compete with the mutant male’s sperm for fertilisations is then 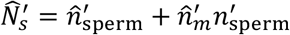, where

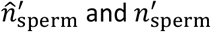 denote the number of sperm that a mutant and a resident male contribute to the group, respectively. The mutant’s share of fertilisation is therefore 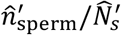, and the total number of fertilisations it achieves is 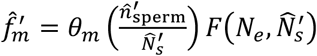. The invasion fitness of the mutant male is thus 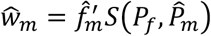. Analogously, the invasion fitness of a mutant female in this scenario is given by 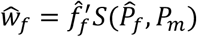, where 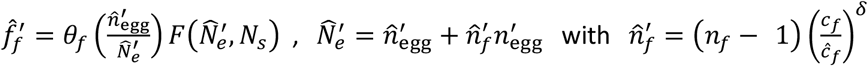.

### Invasion fitness in the presence of uncertain paternity

Our baseline model assumes that individuals of both sexes care only for their own offspring. Extending this baseline model to account for uncertainty in male parentage under scramble competition, we introduce a parameter *η* that represents the degree of uncertain paternity, all else being equal. We assume that females can always identify their own offspring, whereas males cannot discriminate between their own and unrelated offspring. In the presence of uncertain paternity, males may provide care to offspring they do not sire, while some of their own offspring may be cared for by other males. A male’s fitness is therefore determined by two components: (i) the expected number of surviving offspring he sires and cares for himself, and (ii) the expected number of surviving offspring he sires but that are cared for by other males. In a resident population, all males are identical, and thus fitness is independent of *η*. For a rare mutant male, however, invasion fitness depends on both components and is given by 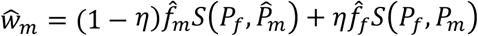 under scramble competition (and 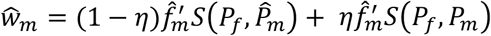 under contest competition), where the first term represents the number of surviving offspring sired and cared for by the mutant, and the second term represents the number of surviving offspring sired by the mutant but cared for by resident males.

### Invasion fitness with depreciable parental care

In the non-depreciable case, invasion fitness is straightforward for all the above scenarios. For a rare mutant male, each of its offspring receives care of 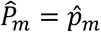 from the mutant male and *P*_*f*_ = *p*_*f*_ from the resident female. A rare female mutant is analogous. Under depreciable care, however, parental investment must be divided among offspring. For a rare mutant male under contests competition, the realised total number of offspring produced is 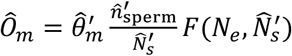, where 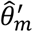 is the mean number of mating groups entered by the mutant conditional on entering at least one group, and the second term gives the number of eggs fertilised by the mutant male within each group. Each offspring therefore receives a fraction 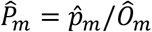 of resources from the mutant male for its survival. Resident females distribute their care across all offspring they produce. Each resident female enters 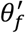 groups (conditional on entering at least one), of which 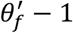 do not contain the mutant male and one contains the mutant male. In groups without the mutant male, a resident female fertilises 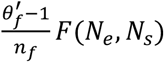eggs across groups, whereas in the group containing the mutant male, it fertilises ^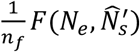^ eggs. As a result, the total number of offspring produced by a resident female 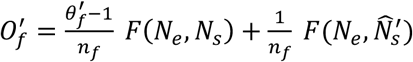, such that each offspring of the mutant male receives a fraction 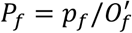 of resources from the resident females determining its survival. The equations for mutant females are analogous. Note that the same calculation applies under scramble competition, with differences arising in total fertilisation *F*, because all eggs and sperm within a group are effectively available for fertilisation, as shown above. Moreover, such depreciable care can also be incorporated into the invasion fitness analysis in the presence of uncertain paternity, with the amount of care received by each offspring of a mutant calculated as described here.

For each scenario, selection gradients, indicating the strength and direction of selection, were approximated numerically for male and female allocation strategies. This allowed us to analyse evolutionary trajectories and identify evolutionary endpoints (see the Mathematica code in the Supplementary Materials for details).

## Data availability

This study is based on mathematical analysis, and no original empirical data were generated. All results can be reproduced using the equations presented in the article and the Mathematica code provided in the Supplementary Materials.

## Acknowledgements

X.L. and J.M.H. acknowledge funding from the Bundesministerium für Bildung und Forschung, Germany, and the Deutsche Forschungsgemeinschaft (DFG, German Research Foundation) under project number 456626331.

## Author Contributions

X.L. and J.M.H. conceived the ideas and designed the model. X.L. performed the mathematical analysis and wrote the manuscript. Both authors interpreted the results and revised the manuscript.

## Preprint Servers

This manuscript is deposited as a preprint on *bioRxiv*. The copyright holders of this preprint are the authors, who have granted *bioRxiv* a license to display the preprint in perpetuity.

## References

1. Darwin, C. The Descent of Man, and Selection in Relation to Sex. (John Murray, 1871).

2. Bateman, A. J. Intra-sexual selection in Drosophila. Heredity 349–368 (1948).

3. Trivers, R. L. Parental investment and sexual selection. in Sexual Selection and the Descent of Man 1871–1971 (eds Houck, L. D. & Drickamer, L. C. 795–838 (University of Chicago Press, 1972).

4. Clutton-Brock, T. H. The Evolution of Parental Care. (Princeton University Press, 1991).

5. Dawkins, R. & Carlisle, T. R. Parental investment, mate desertion and a fallacy. Nature 262, 131–133 (1976).

6. Alonzo, S. H. Social and coevolutionary feedbacks between mating and parental investment. Trends Ecol. Evol. 25, 99–108 (2010).

7. Long, X. & Weissing, F. J. Transient polymorphisms in parental care strategies drive divergence of sex roles. Nat. Commun. 14, 6805 (2023).

8. Blumer, L. S. Male parental care in the bony fishes. Q. Rev. Biol. 54, 149–161 (1979).

9. Balshine, S. Patterns of parental care in vertebrates. in The Evolution of Parental Care (eds Royle, N. J., Smiseth, P. T. & Kölliker, M.) 62–80 (Oxford University Press, 2012).

10. Furness, A. I. & Capellini, I. The evolution of parental care diversity in amphibians. Nat. Commun. 10, 4709 (2019).

11. Tang-Martinez, Z. Bateman’s principles: original experiment and modern data for and against. in Encyclopedia of Animal Behavior (eds Breed, M. D. & Moore, J.) 166–176 (Oxford University Press, 2010).

12. Janicke, T., Häderer, I. K., Lajeunesse, M. J. & Anthes, N. Darwinian sex roles confirmed across the animal kingdom. Sci. Adv. 2, e1500983 (2016).

13. Queller, D. C. Why do females care more than males? Proc. R. Soc. Lond. B Biol. Sci. 264, 1555–1557 (1997).

14. Kokko, H. & Jennions, M. D. Parental investment, sexual selection and sex ratios. J. Evol. Biol. 21, 919–948 (2008).

15. McNamara, J. M. & Wolf, M. Sexual conflict over parental care promotes the evolution of sex differences in care and the ability to care. Proc. R. Soc. B Biol. Sci. 282, 20142752 (2015).

16. de Vries, C. & Lehtonen, J. Sex-specific assumptions and their importance in models of sexual selection. Trends Ecol. Evol. 38, 927–935 (2023).

17. Fromhage, L. & Jennions, M. D. Coevolution of parental investment and sexually selected traits drives sex-role divergence. Nat. Commun. 7, (2016).

18. Iyer, P., Shukla, A., Jadhav, V. & Sahoo, B. K. Anisogamy selects for male-biased care in self-consistent games with synchronous matings. Evolution 74, 1018–1032 (2020).

19. Lehtonen, J., Parker, G. A. & Schärer, L. Why anisogamy drives ancestral sex roles. Evolution 70, 1129–1135 (2016).

20. Lehtonen, J. Bateman gradients from first principles. Nat. Commun. 13, (2022).

21. Gowaty, P. A. & Hubbell, S. P. Chance, time allocation, and the evolution of adaptively flexible sex role behavior. Integr. Comp. Biol. 45, 931–944 (2005).

22. Gowaty, P. A. & Hubbell, S. P. Reproductive decisions under ecological constraints: it’s about time. Proc. Natl. Acad. Sci. 106, 10017–10024 (2009).

23. Schärer, L., Rowe, L. & Arnqvist, G. Anisogamy, chance and the evolution of sex roles. Trends Ecol. Evol. 27, 260–264 (2012).

24. Ah-King, M. On anisogamy and the evolution of ‘sex roles’.Trends Ecol. Evol. 28, 1–2 (2013).

25. Hubbell, S. P. & Johnson, L. K. Environmental variance in lifetime mating success, mate choice, and sexual selection. Am. Nat. 130, 91–112 (1987).

26. Janicke, T. Sexual selection: competition for resources provided by mating partners. Behav. Ecol. 35, arae016 (2024).

27. Herberstein, M. E., Painting, C. J. & Holwell, G. I. Scramble competition polygyny in terrestrial arthropods. in Advances in the Study of Behavior (eds Naguib, M. et al.) 237–295 (Elsevier Academic Press, 2017).

28. Boisseau, R. P. et al. Multi-modal locomotor costs favor smaller males in a sexually dimorphic leaf-mimicking insect. BMC Ecol. Evol. 22, (2022).

29. Wells, K. D. The Ecology and Behavior of Amphibians. (University of Chicago Press, 2007).

30. Lane, J. E., Boutin, S., Speakman, J. R. & Humphries, M. M. Energetic costs of male reproduction in a scramble competition mating system. J. Anim. Ecol. 79, 27–34 (2010).

31. Thornhill, R. & Alcock, J. The Evolution of Insect Mating Systems. (Harvard University Press, 1983).

32. Andersson, M. Sexual Selection. (Princeton University Press, 1994).

33. Smith, C. & Wootton, R. J. The costs of parental care in teleost fishes. Rev. Fish Biol. Fish. 5, 7–22 (1995).

34. Mank, J. E., Promislow, D. E. L. & Avise, J. C. Phylogenetic perspectives in the evolution of parental care in ray-finned fishes. Evolution 59, 1570–1578 (2005).

35. Lehtonen, J. & Parker, G. A. The correlation between anisogamy and sexual selection intensity—the broad theoretical predictions. Evol. Lett. 8, 749–755 (2024).

36. Mokos, J., Scheuring, I., Liker, A., Freckleton, R. P. & Székely, T. Degree of anisogamy is unrelated to the intensity of sexual selection. Sci. Rep. 11, (2021).

37. Schärer, L. Tests of sex allocation theory in simultaneously hermaphroditic animals. Evolution 63, 1377–1405 (2009).

38. Lehtonen, J. & Kokko, H. Two roads to two sexes: unifying gamete competition and gamete limitation in a single model of anisogamy evolution. Behav. Ecol. Sociobiol. 65, 445–459 (2011).

39. Henshaw, J. M., Jones, A. G. & Schärer, L. Anisogamy explains why males benefit more from additional matings. Nat. Commun. 13, 3893 (2022).

40. Santos, E. S. A., Bueno, P. P., Gilbert, J. D. J. & Machado, G. Macroecology of parental care in arthropods: higher mortality risk leads to higher benefits of offspring protection in tropical climates. Biol. Rev. 92, 1688–1701 (2017).

41. Goldberg, R. L., Downing, P. A., Griffin, A. S. & Green, J. P. The costs and benefits of paternal care in fish: a meta-analysis. Proc. R. Soc. B Biol. Sci. 287, 20201759 (2020).

42. Machado, G. & Macedo-Rego, R. C. Benefits and costs of female and male care in amphibians: a meta-analytical approach. Proc. R. Soc. B Biol. Sci. 290, 20231759 (2023).

43. Griffith, S. C., Owens, I. P. F. & Thuman, K. A. Extra pair paternity in birds: a review of interspecific variation and adaptive function. Mol. Ecol. 11, 2195–2212 (2002).

44. Cohas, A. & Allainé, D. Social structure influences extra-pair paternity in socially monogamous mammals. Biol. Lett. 5, 313–316 (2009).

45. Höglund, J. & Alatalo, R. V. Leks. (Princeton University Press, 1995).

46. Wiley, R. H. Lekking in birds and mammals: behavioral and evolutionary issues. in Advances in the Study of Behavior (eds Slater, P. J. B., Rosenblatt, J. S., Beer, C. & Milinski, M.) 201–291 (Academic Press, 1991).

47. Fiske, P., Rintamäki, P. T. & Karvonen, E. Mating success in lekking males: a meta-analysis. Behav. Ecol. 9, 328–338 (1998).

48. Sherman, P. W. Birds of a feather lek together. Nature 401, 119–120 (1999).

49. Hayward, A. & Gillooly, J. F. The cost of sex: quantifying energetic investment in gamete production by males and females. Plos One 6, e16557 (2011).

50. Emlen, S. T. & Oring, L. W. Ecology, sexual selection, and the evolution of mating systems. Science 197, 215–223 (1977).

51. Kokko, H. & Rankin, D. J. Lonely hearts or sex in the city? Density-dependent effects in mating systems. Philos. Trans. R. Soc. B Biol. Sci. 361, 319–334 (2006).

52. Alonzo, S. H. & Klug, H. Paternity, maternity, and parental care. in The Evolution of Parental Care (eds Royle, N. J., Smiseth, P. T. & Kölliker, M.) 189–205 (Oxford University Press, 2012).

53. Mayer, C. & Pasinelli, G. New support for an old hypothesis: density affects extra-pair paternity. Ecol. Evol. 3, 694–705 (2013).

54. Maldonado-Chaparro, A. A., Montiglio, P.-O., Forstmeier, W., Kempenaers, B. & Farine, D. R. Linking the fine-scale social environment to mating decisions: a future direction for the study of extra-pair paternity. Biol. Rev. 93, 1558–1577 (2018).

55. Long, X., Lehtonen, J. & Henshaw, J. M. Quantifying feedback among traits in coevolutionary models. Evolution 79, 2597–2611 (2025).

56. Summers, K., Weigt, L. A., Boag, P. & Bermingham, E. The evolution of female parental care in poison frogs of the genus Dendrobates: evidence from mitochondrial DNA sequences. Herpetologica 55, 254–270 (1999).

57. Gross, M. R. & Sargent, R. C. The evolution of male and female parental care in fishes. Am. Zool. 25, 807–822 (1985).

58. Reynolds, J. D., Goodwin, N. B. & Freckleton, R. P. Evolutionary transitions in parental care and live bearing in vertebrates. Philos. Trans. R. Soc. B Biol. Sci. 357, 269–281 (2002).

59. Wesolowski, T. On the origin of parental care and the early evolution of male and female parental roles in birds. Am. Nat. 143, 39–58 (1994).

60. Wesołowski, T. The origin of parental care in birds: a reassessment. Behav. Ecol. 15, 520–523 (2004).

61. Henshaw, J. M., Fromhage, L. & Jones, A. G. Sex roles and the evolution of parental care specialization. Proc. R. Soc. B Biol. Sci. 286, 20191312 (2019).

62. Long, X. et al. Does ecology and life history predict parental cooperation in birds? A comparative analysis. Behav. Ecol. Sociobiol. 76, 92 (2022).

63. Henshaw, J. M., Fromhage, L. & Jones, A. G. The evolution of mating preferences for genetic attractiveness and quality in the presence of sensory bias. Proc. Natl. Acad. Sci. 119, e2206262119 (2022).

64. Kuijper, B., Pen, I. & Weissing, F. J. A guide to sexual selection theory. Annu. Rev. Ecol. Evol. Syst. 43, 287–311 (2012).

65. Gross, M. R. Alternative reproductive strategies and tactics: diversity within sexes. Trends Ecol. Evol. 11, 92–98 (1996).

66. Taborsky, M., Oliveira, R. F. & Brockmann, H. J. The evolution of alternative reproductive tactics: concepts and questions. in Alternative Reproductive Tactics: An Integrative Approach (eds Oliveira, R. F., Taborsky, M. & Brockmann, H. J.) 25–51 (Cambridge University Press, 2008).

67. Wang, D. et al. Female alternative reproductive tactics: diversity and drivers. Trends Ecol. Evol. 39, 937–946 (2024).

68. Kokko, H. & Mappes, J. Sexual selection when fertilization is not guaranteed. Evolution 59, 1876–1885 (2005).

