## Supplementary information for "Anisogamy drives diverse patterns of sex-role divergence"

[2]Present address: Department of Biology, University of Konstanz, Universitaetsstrasse 10, 78464 Constance, Germany

[*]To whom correspondence should be addressed.

This PDF file includes:

Description of the derivation of the fertilisation function in our model.

Supplementary Figure 1. Effect of fertilisation efficiency on total fertilisation

Supplementary Figure 2. Fertilisation efficiency affects sex role evolution.

Supplementary Figure 3. Effect of local group size on sex role evolution under scramble competition.

Supplementary Figure 4. Offspring baseline survival influences sex role evolution.

Supplementary Figure 5. Effect of local group size on sex role evolution under contest competition.

Supplementary Figure 6. Anisogamy favours typical sex roles across initial conditions.

Supplementary Figure 7. Depreciable care reshapes allocation strategies under scramble competition.

Supplementary Figure 8. Effect of local group size on sex role evolution under scramble competition with depreciable care.

Supplementary Figure 9. Resource allocation under scramble competition with depreciable post-zygotic care and uncertain paternity.

Supplementary Figure 10. Anisogamy alone drives male-biased care.

Description of the feedback analysis quantifying the causal pathway from anisogamy to male care.

Supplementary Figure 11. Evolutionary trajectories of resource allocation without mating competition.

Supplementary Figure 12. Feedback pathways linking anisogamy to selection on female traits.

Supplementary Figure 13. Feedback pathways linking anisogamy to selection on male traits.

Table 1. Overview of notation in the main text.

SI references

The Supplementary Information is organised as follows. We first describe the derivation of the fertilisation function and show how the fertilisation efficiency parameter affects total fertilisation (Supplementary Fig. 1). We then systematically investigate how key ecological parameters affect sex-role evolution (Supplementary Figs. 2-9). To clarify the causal link from anisogamy to post-zygotic care, we analyse a simplified version of the model in which resources are allocated only between gamete production and parental care (Supplementary Fig. 10), then apply the feedback analysis framework to quantitatively decompose the causal pathways involved and present the corresponding results (Supplementary Figs. 11-13). We conclude with an overview of the model notation used throughout the main text (Table 1), followed by the references for the Supplementary Information.

**Gamete fertilisation within local mating groups.** As described in the Methods of the main text, resident females and males produce a total of $N_e$ eggs and $N_s$ sperm, respectively. However, the number of gametes available for fertilisation is dynamic, as gametes experience instantaneous mortality and fertilisation after being produced.

Following previous studies on the fertilisation dynamics of gametes produced by two mating types[1,2], we assume that the breeding season is sufficiently long for gamete dynamics to reach a steady state, where the influx of newly produced gametes balances losses due to mortality and fertilisation. Rather than a literal representation of the biology, this assumption should be viewed as a pragmatic tool to obtain a fertilisation function with sensible properties. Let $n_e$ and $n_s$ denote the numbers of eggs and sperm available for fertilisation at any given time. Their dynamics are then given by

$$\frac{dn_e}{dt} = N_e - \mu_e n_e - \gamma n_e n_s, \quad \text{(S1a)}$$

$$\frac{dn_s}{dt} = N_s - \mu_s n_s - \gamma n_e n_s, \quad \text{(S1b)}$$

where $\mu_e$ and $\mu_s$ are the instantaneous mortality rates of eggs and sperm, respectively, and $\gamma$ represents fertilisation efficiency.

Setting equations $(S1a)$ and $(S1b)$ to zero yields the equilibrium numbers of eggs and sperm available for fertilisation. Substituting these equilibrium values into the expression for the total fertilisation rate, $\gamma n_e n_s$, gives

$$F(N_e, N_s) = \frac{\gamma(N_e+N_s)+\mu_e\mu_s-\sqrt{\gamma^2(N_e-N_s)^2+2\gamma(N_e+N_s)\mu_e\mu_s+\mu_e^2\mu_s^2}}{2\gamma}. \quad \text{(S2)}$$

For simplicity, we assume equal gamete mortality rates ($\mu_e = \mu_s$) and set them to 1 without loss of generality. By doing so, the fertilisation function simplifies to

$$F(N_e, N_s) = \frac{1+\gamma(N_e+N_s)-\sqrt{1+\gamma^2(N_e-N_s)^2+2\gamma(N_e+N_s)}}{2\gamma}. \qquad \text{(S3)}$$

This fertilisation function has several desirable properties for our model. First, it is symmetric in gamete numbers, such that $F(N_e, N_s) = F(N_s, N_e)$. Second, it provides a smooth transition from zero fertilisation at low efficiency ($\gamma \to 0$) to complete fertilisation at high efficiency, where the number of fertilised eggs approaches $\min(N_e, N_s)$. In particular, as fertilisation efficiency increases ($\gamma \to \infty$), the function approaches $N_e$ when sperm are not limiting (i.e., $N_s \geq N_e$), including both $N_s \gg N_e$and $N_s = N_e$.

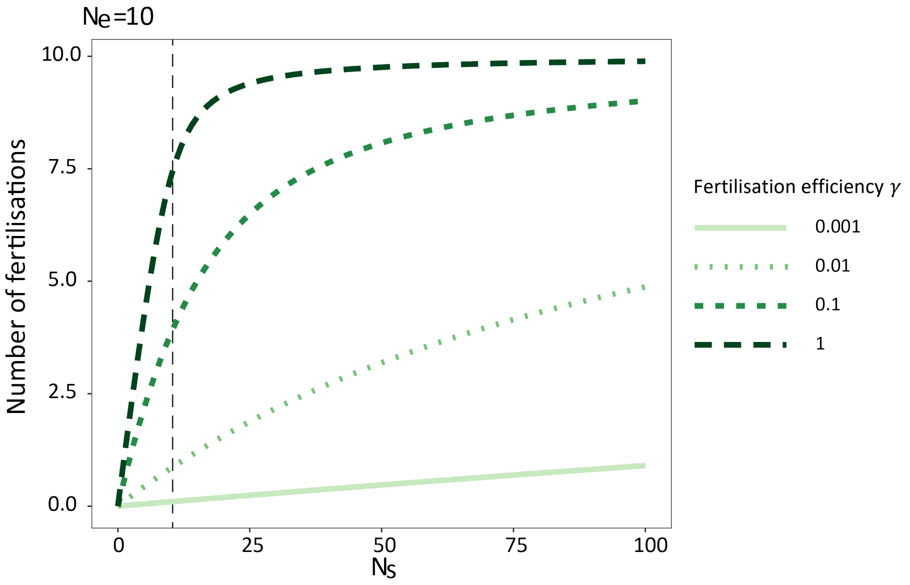


**Supplementary Figure 1. Effect of fertilisation efficiency on total fertilisation.** The figure shows how fertilisation efficiency $\gamma$ influences the total number of fertilisations. Egg number is fixed at $N_e = 10$, while sperm number ($N_s$) ranges from 0 to 100. Increasing fertilisation efficiency leads to a higher number of fertilised eggs for a given sperm abundance. Because eggs are typically larger than sperm, a given resource budget generally produces more sperm than eggs. Thus, scenarios in which sperm number is lower than egg number (left of the dashed vertical line at $N_s = 10$) are expected to be relatively rare. In most of cases (to the right of the dashed vertical line), fertilisation becomes less limited by sperm availability as fertilisation efficiency increases, and the number of fertilised eggs approaches the total number of eggs present when the fertilisation efficiency is sufficiently high. Throughout the manuscript, the default value of fertilisation efficiency is $\gamma = 1$, under which most eggs are fertilised when sperm are moderately abundant.

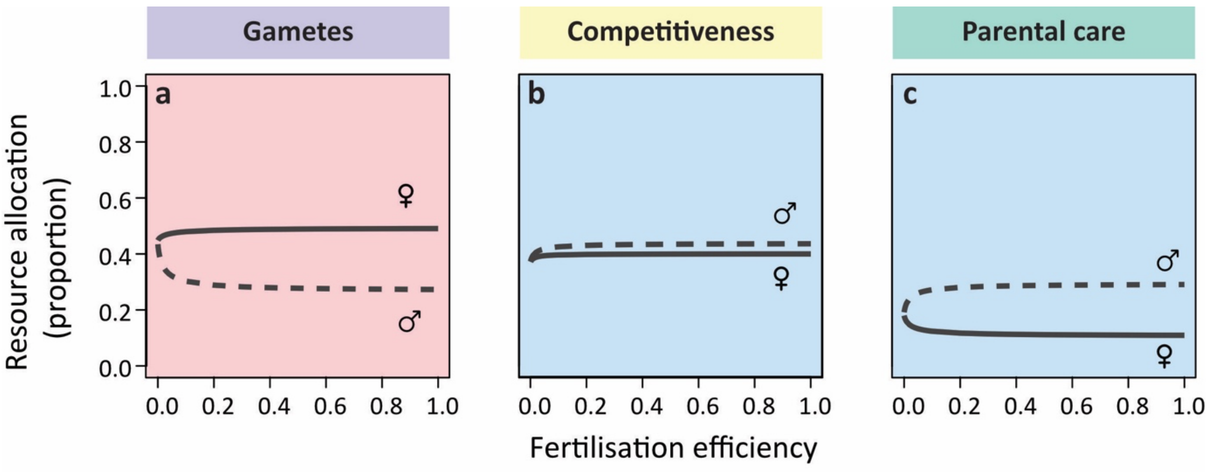


**Supplementary Figure 2. Fertilisation efficiency affects sex role evolution.** The graph shows how fertilisation efficiency influences the link between anisogamy and subsequent sex-role evolution under scramble competition. Here, all model assumptions and parameter settings are identical to those in Fig. 2 of the main text with egg size $m_f = 0.1$ and sperm size $m_m = 0.01$, except that fertilisation efficiency is varied. When fertilisation efficiency decreases, saturation in the relationship between sperm number and fertilisation success is reduced (Supplementary Fig. 1), thereby linearising fitness returns on sperm production[3–5]. Males consequently redirect resources away from mating competition and parental care and towards sperm production, while females shift resources from the other two traits to increase investment in post-zygotic care in response to reduced male care. Note that the default value used throughout the manuscript is $\gamma = 1$.

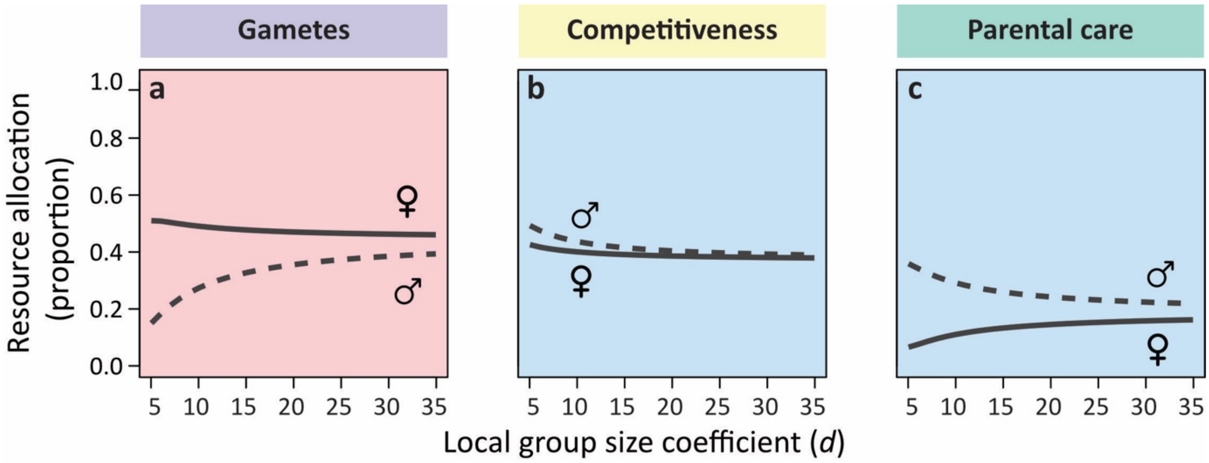


**Supplementary Figure 3. Effect of local group size on sex role evolution under scramble competition.** This graph shows how expected local mating group size influences the pathway linking anisogamy to secondary sex-role evolution under scramble competition. Again, we retain the model assumptions and parameter settings from Fig. 2 of the main text, with egg size $m_f = 0.1$ and sperm size $m_m = 0.01$, while varying expected local mating-group size $d$. Under scramble competition, the parameter $d$ is not directly equal to the realised group size, because the absolute competitiveness determines how many local groups an individual can enter, which also affects the realised number of males and females present in each group. However, for a given level of competitiveness, larger values of $d$ correspond to larger realised group sizes. As $d$ increases, individuals therefore become more aggregated within local mating groups, reducing selection for mate-encounter traits in both sexes because potential mates are easier to encounter locally. At the same time, larger groups bring more males into competition for a limited number of eggs, thereby intensifying sperm competition. Stronger sperm competition makes fitness returns on sperm production more nearly linear, favouring increased male allocation to sperm production at the expense of both mating competition and parental care. In response to reduced male care, females are favoured to increase investment in parental care, which comes at the expense of egg production. This reduction in egg availability can further strengthen sperm competition within local groups. As a result, the degree of secondary sex-role divergence decreases as expected local group size increases. Note that the default value used throughout the manuscript is $d = 10$.

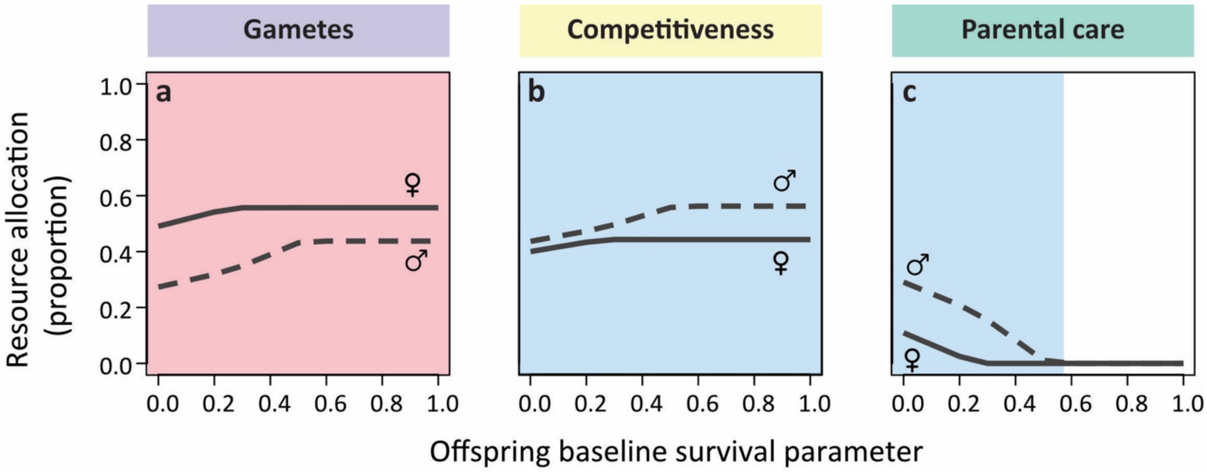


**Supplementary Figure 4. Offspring baseline survival influences sex role evolution.** In our model, offspring survival is determined by baseline survival and the total care received from both parents (see Methods in the main text). Whereas Fig. 2 assumes that offspring cannot survive without any care, here we vary the baseline survival parameter $b$ to examine how care dependence affects sex-role divergence. In the absence of parental care, offspring survival is given by $S(0,0) = 1 - \mathrm{e}^{-b}$; thus, as $b$ increases, offspring survival becomes less dependent on parental care. Accordingly, increasing the baseline survival parameter reduces parental investment by both sexes, with female care declining to zero first. This leads to a transition from male-biased care to male-only care, and ultimately to no parental care when baseline survival is sufficiently high. The resources released from post-zygotic care are then redirected towards gamete production and mating competition. Females mainly increase egg production, while retaining enough investment in mating competition to encounter males. Males instead invest more heavily in scramble competition than in sperm production, because realised local group size remains small and local sperm competition is therefore strong.

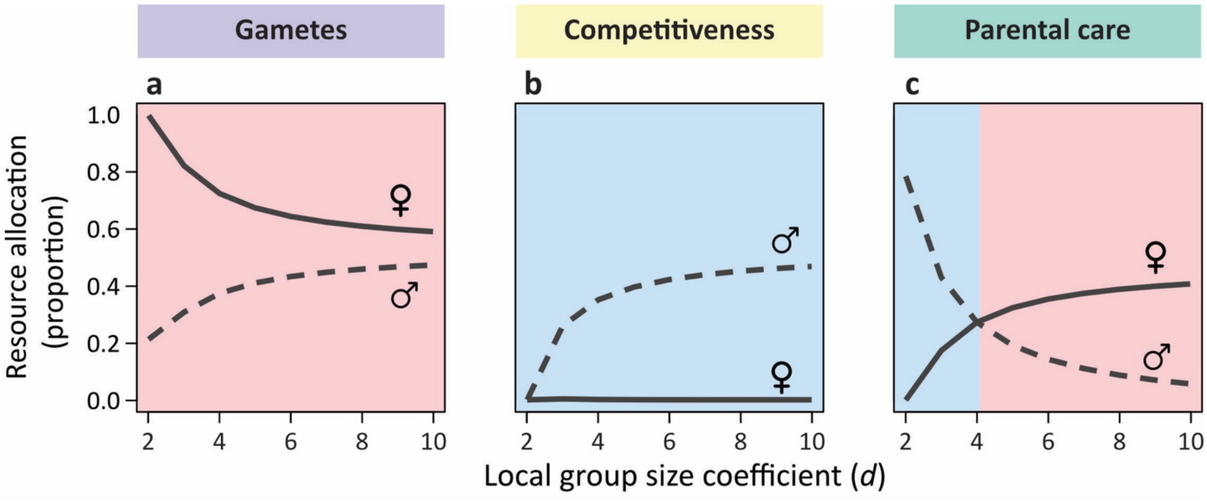


**Supplementary Figure 5. Effect of local group size on sex role evolution under contest competition.** This graph shows how expected local mating group size alters opportunities for mate monopolisation, and thus reshapes sex-role divergence under contest competition. Here, we retain the model assumptions and parameter settings from Fig. 4 of the main text, with egg size $m_f = 0.1$ and sperm size $m_m = 0.01$, while varying expected local mating-group size $d$. Under contest competition, the parameter $d$ directly corresponds to realised group size, because individuals are assumed to enter a fixed number of groups. Accordingly, small local groups provide little opportunity to monopolise multiple mates, so contest competition remains weak or even absent. As a result, females allocate more resources to egg production, whereas males invest more in parental care. As local groups become larger, opportunities for mate monopolisation increase, intensifying contest competition, particularly among males whose reproductive success is limited by access to eggs. Females consequently increase parental care at the expense of egg production.

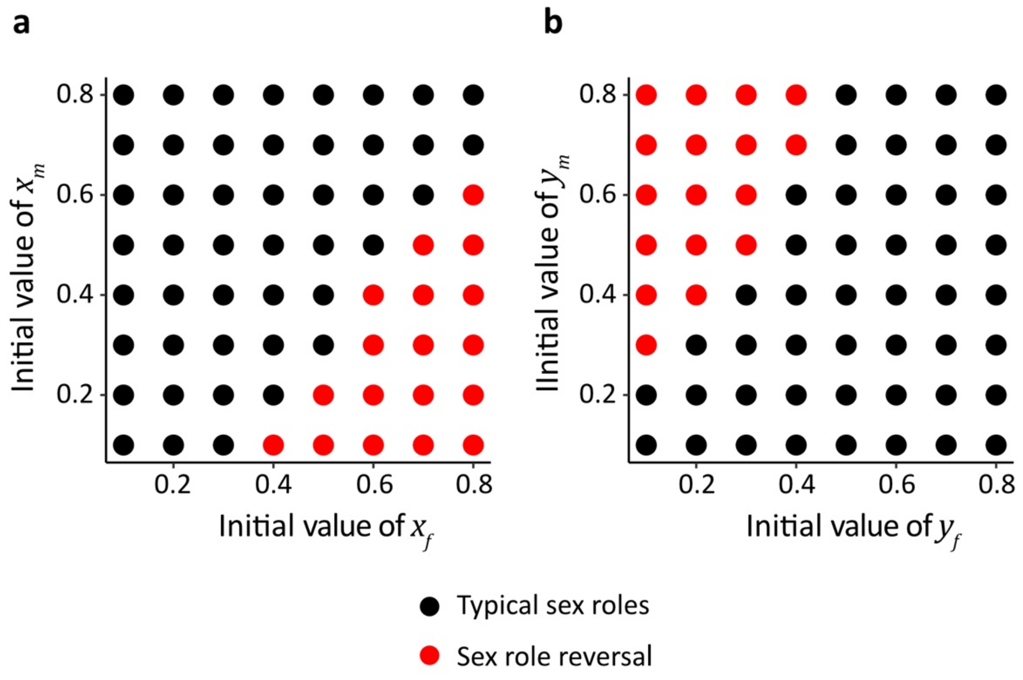


**Supplementary Figure 6. Anisogamy favours typical sex roles across initial conditions.** As shown in Fig. 6 of the main text, under contest competition with depreciable parental care, populations converge to two alternative resource-allocation strategies depending on initial conditions: one characterised by typical sex roles, with competing males and caring females, and the other by sex-role reversal, with competing females and caring males. However, because anisogamy constrains the evolutionary outcomes, most initial conditions lead populations towards typical sex roles. Sex-role reversal evolves only **(a)** when females initially exhibit greater competitiveness than males, or **(b)** when males initially provide substantially more care than females. Thus, the dynamics are not fully symmetric across initial conditions. Here, all model assumptions and parameter settings are identical to those in Fig. 6 of the main text, with egg size $m_f = 0.1$ and sperm size $m_m = 0.01$, but a wider range of initial values for the evolving resource allocation traits is explored. Black dots represent the typical sex-role equilibrium, whereas red dots represent the complete sex-role reversal equilibrium.

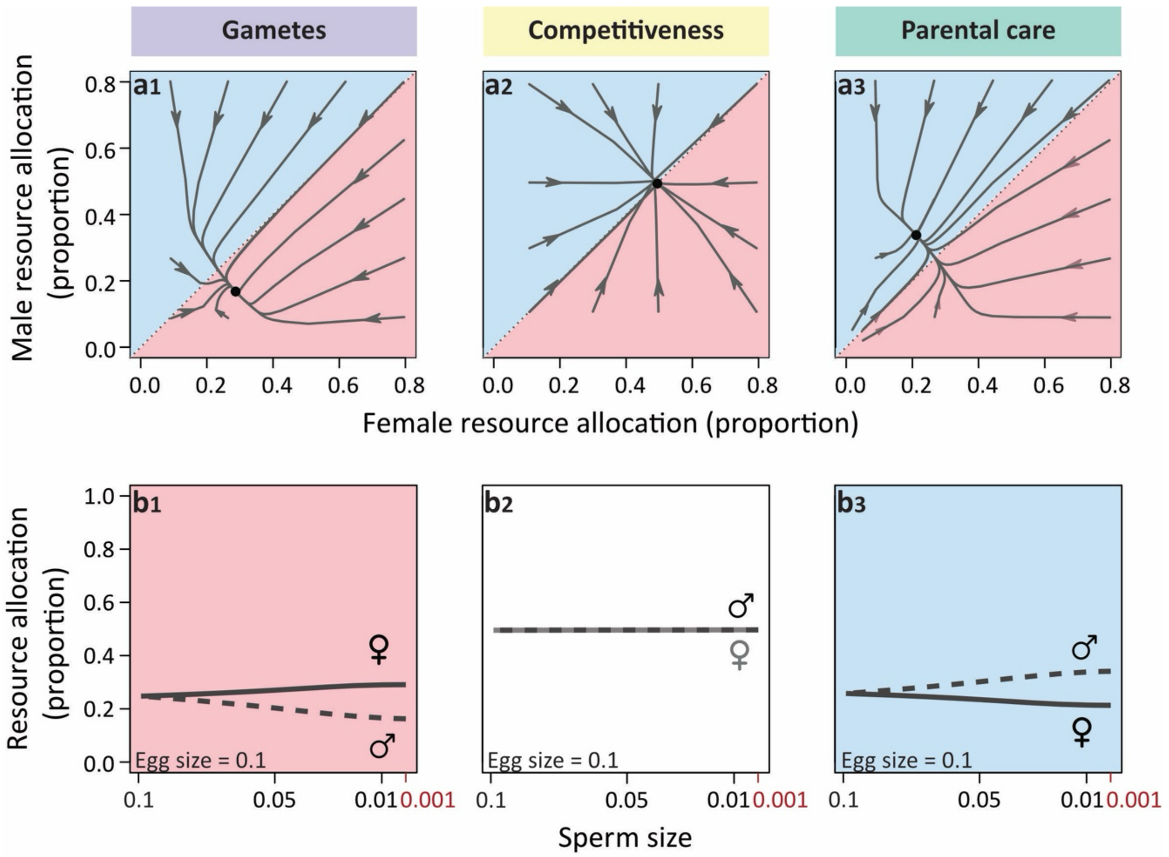


**Supplementary Figure 7. Depreciable care reshapes allocation strategies under scramble competition.** The figure shows how anisogamy drives secondary sex-role divergence under scramble competition when post-zygotic care is depreciable. All assumptions and parameter settings are as in Fig. 2 of the main text, except that resources allocated to parental care are divided equally among all offspring produced by an individual parent (see Methods in the main text for details). The model yields qualitatively similar patterns, with female-biased investment in gamete production and male-biased investment in post-zygotic care. However, several important differences emerge. First, total investment in egg production is strongly reduced compared to the case of non-depreciable care (**a**$_1$,**b**$_1$), leading to substantially smaller clutch sizes. Second, females become approximately as competitive as males (**a**$_2$,**b**$_2$). This is because males provide more parental care than females (**a**$_3$,**b**$_3$), male care becomes a limiting resource for females in this scenario. Consequently, females compete for access to caring males, and therefore able to provide higher per-offspring investment. Thus, depreciable care introduces an additional dimension of mating competition by making per-offspring parental investment a limiting and valuable resource. As a result, mating competition becomes two-sided: males compete for fertilization opportunities over a limited number of eggs, while females compete to mate with multiple caring males, effectively distributing offspring across partners to enhance per-offspring care. Increasing expected local group size reduces sex differences in resource allocation across the three reproductive activities (Supplementary Fig. 8), whereas increasing paternity uncertainty shifts the pattern towards typical sex roles, with greater male investment in competition and greater female investment in care, alongside smaller clutch sizes (Supplementary Fig. 9).

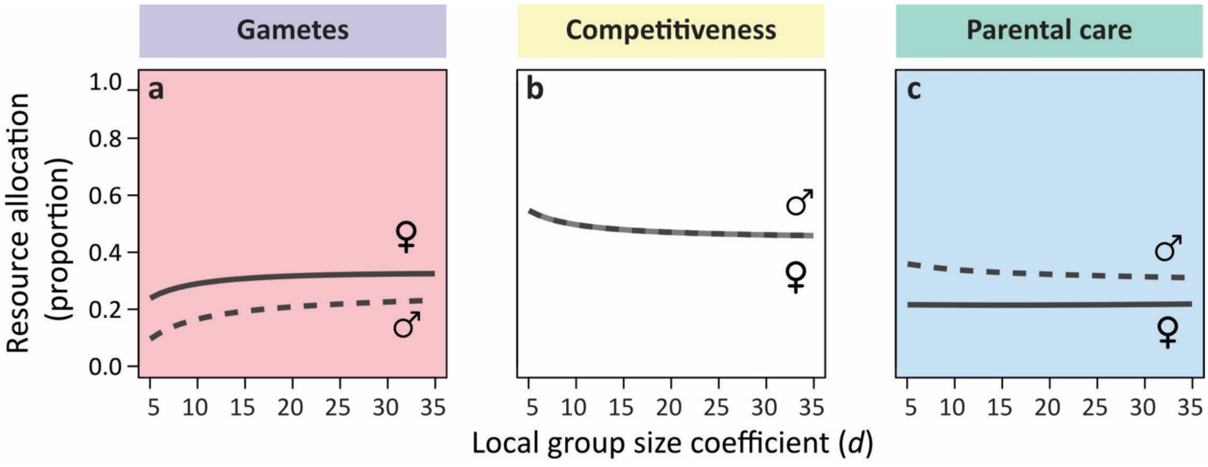


**Supplementary Figure 8. Effect of local group size on sex role evolution under scramble competition with depreciable care.** Females and males allocate resources to (**a**) gamete production, (**b**) mating competition, and (**c**) post-zygotic parental care under scramble competition with depreciable parental care and varying expected group size. Here, increasing expected local group size reduces the need for active mate searching or long-distance signalling in both sexes, as opposite sex individuals become less limiting. Consequently, both males and females decrease investment in scramble competition (b). However, as the number of males per group increases, sperm competition intensifies, leading males to allocate more resources to sperm production and less to parental care (a,c). In contrast, females do not compensate for the reduction in paternal care, but instead increase investment in egg production. This reflects a quality–quantity trade-off in offspring production: producing more offspring, each receiving less total care, is favoured over producing fewer offspring that each receive more care. All model assumptions and parameter settings are as in Supplementary Fig. 6a, except that the expected group size parameter $d$ is varied.

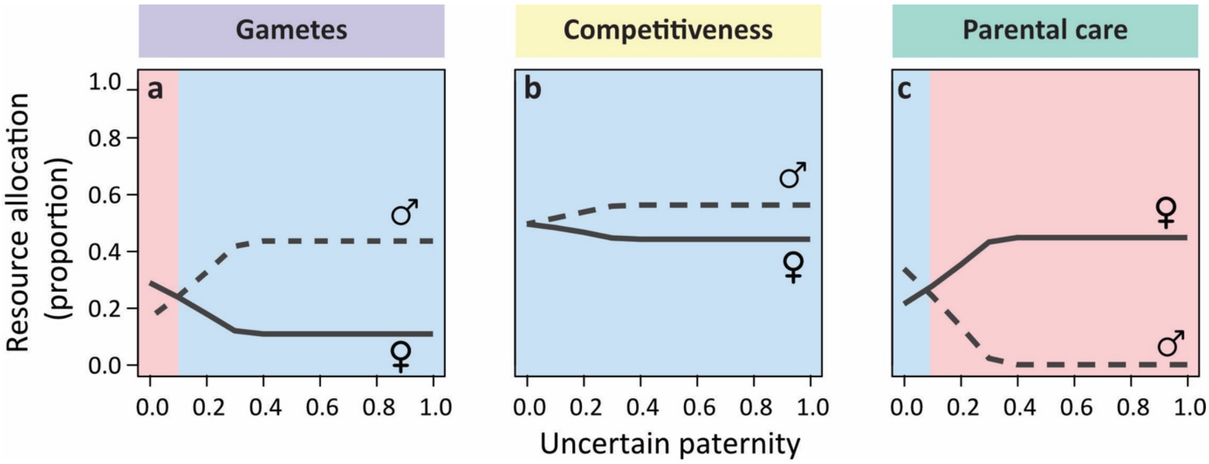


**Supplementary Figure 9. Resource allocation under scramble competition with depreciable post-zygotic care and uncertain paternity.** All model assumptions and parameter settings are as in Supplementary Fig. 6a, except that uncertain paternity is incorporated, as in Fig. 3 of the main text. Increasing uncertain paternity produces qualitatively similar effects to those observed under non-depreciable care (cf. Fig. 3), but the response is markedly more sensitive here. Even low levels of uncertain paternity substantially shift the equilibrium pattern relative to the non-depreciable case. Under depreciable care, females produce fewer eggs than in the non-depreciable scenario (**a**) and allocate more resources to maternal care, which is divided among offspring (**c**). In addition, sex differences in mating competition re-emerge as uncertain paternity increases (**b**). Since males reduce paternal care and eventually provide no care, the benefit for females to compete for access to males diminishes. Consequently, females reduce investment in mating competition to the level needed to ensure access enough local groups, as observed in the non-depreciable case.

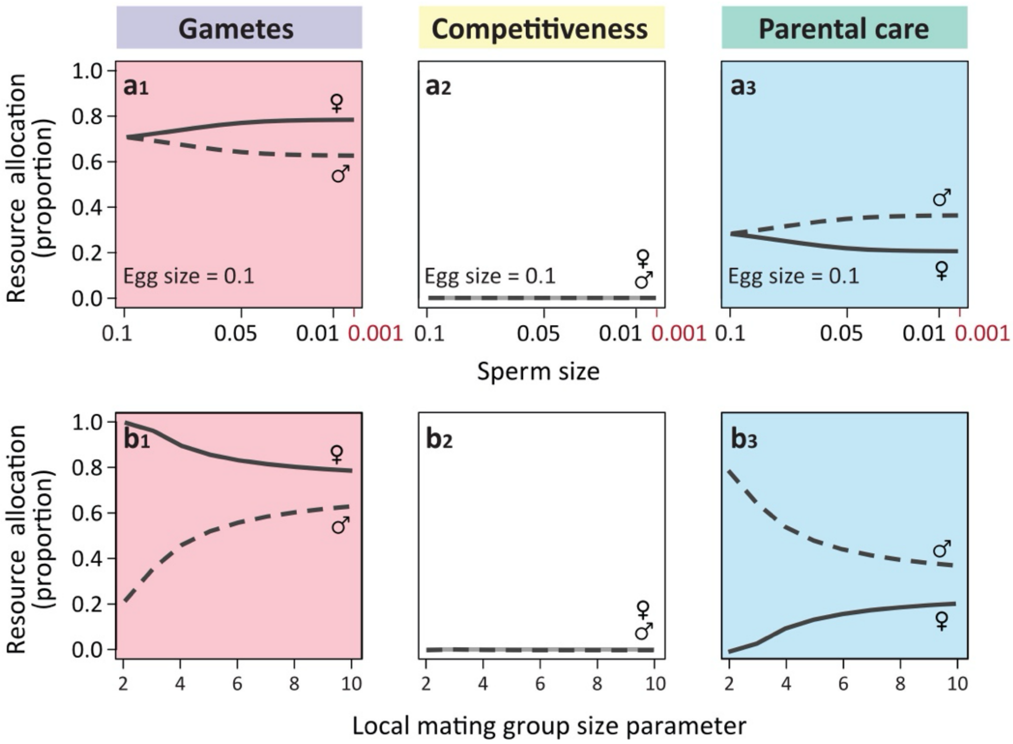


**Supplementary Figure 10. Anisogamy alone can drive male-biased care.** The panels depict resource allocation strategies when males and females enter a fixed number of local mating groups, and mating occurs randomly within each group. Under these conditions, individuals do not invest in any forms of pre-mating competition **($b_1$,$b_2$).** When the fertilisation rate is high, egg production is strongly linked to fitness because additional eggs are very likely to be fertilised. Females are therefore selected to allocate more resources to egg production **($a_1$,$a_2$)**, shifting resources away from post-zygotic care (**$c_1$**,**$c_2$**) This reduction in female care, in turn, favours increased male care because offspring survival depends on the total post-zygotic care provided by both parents. (**a**) These sex differences in resource allocation are amplified as the anisogamy ratio increases. Moreover, (**b**) as local group size increases, sperm competition becomes stronger, favouring increased male allocation to sperm production and reduced allocation to parental care. As male care declines, females are selected to increase their own allocation to care. Overall, however, males still provide more post-zygotic care than females. Thus, this analysis shows that, in the absence of pre-mating competition, anisogamy is directly linked to male-biased rather than female-biased post-zygotic care. This contrasts with Trivers' classical argument: when the only initial sex difference is gamete size, females do not necessarily evolve to provide most care. Instead, because females produce fewer but larger gametes, eggs have high fitness value when the fertilisation rate is high, favouring increased female investment in egg production. This leaves fewer resources available for post-zygotic care, thereby favouring increased male care. The feedback analysis below, together with in Supplementary Figs. 12 and 13, decomposes and quantifies this causal pathway. Key parameter values: $R = 1$, $r = 0.5$, $\delta = 0$. In both panels (a) and (b), the resource allocation variables are initially set to $x_f = y_f = x_m = y_m = 0.1$. In panel (a), $d = 10$; in panel (b), $m_f = 0.1$, $m_m = 0.01$.

**Feedback analysis quantifies the causal pathways from anisogamy to male-biased post-zygotic care.** To develop a more comprehensive and quantitative understanding of the causal pathways from anisogamy to the evolution of male-biased post-zygotic parental care shown in Supplementary Fig. 10, we apply a recently developed framework for feedback analysis among coevolving traits[6]. This framework allows us to examine how evolutionary changes in each trait feed back to shape selection on other traits when multiple traits interact, providing a fine-grained and systematic way to quantify coevolutionary feedback between traits and selection gradients and to tease apart the pathways through which they influence one another in an adaptive-dynamics model[6].

To apply this feedback-analysis framework, and in particular to understand the direct causal link between anisogamy and the subsequent distribution of parental care between males and females, we allow individuals to allocate resources only between gamete production and post-zygotic parental care. Thus, a female with trait values $x_f$ and $y_f$ allocates a proportion $g_f = x_f/(x_f + y_f)$ of her resources to egg production and a proportion $p_f = y_f/(x_f + y_f)$ to post-zygotic parental care. The evolving variables $x_f$ and $y_f$ therefore determine the relative allocation of resources to gamete production and parental care, respectively. Male allocation is analogous (i.e. $g_m = x_m/(x_m + y_m), p_m = y_m/(x_m + y_m)$). Based on their resource allocation strategies, individuals then proceed through the life cycle depicted in Fig. 1b, but here without mating competition. We therefore assume that each individual enters a fixed number of local mating groups, set to one by default, and that mating occurs randomly within each group. All other assumptions follow the baseline model described in the main text (with full certainty of paternity and non-depreciable parental care). Accordingly, the invasion fitness $\hat{w}_f(\hat{x}_f, \hat{y}_f, x_m, y_m)$ and $\hat{w}_m(x_f, y_f, \hat{x}_m, \hat{y}_m)$ for a mutant male and a mutant female, respectively, can be derived as in the main text (see also the Mathematica code in the Supplementary Materials for full details of these fitness expressions and feedback analysis).

Based on these invasion fitness functions, the selection gradients for each resource allocation variable are given by:

$$\beta_{x_f} = \frac{1}{\overline{w}_f} \frac{\partial}{\partial \hat{x}_f} w_f(\hat{x}_f, \hat{y}_f, x_m, y_m)|_{\hat{x}_f = x_f, \hat{y}_f = y_f} \quad (1a)$$

$$\beta_{y_f} = \frac{1}{\overline{w}_f} \frac{\partial}{\partial \hat{y}_f} w_f(\hat{x}_f, \hat{y}_f, x_m, y_m)|_{\hat{x}_f = x_f, \hat{y}_f = y_f} \quad (1b)$$

$$\beta_{x_m} = \frac{1}{\overline{w}_m} \frac{\partial}{\partial \hat{x}_m} w_m(x_f, y_f, \hat{x}_m, \hat{y}_m)|_{\hat{x}_m = x_m, \hat{y}_m = y_m} \quad (1c)$$

$$\beta_{y_m} = \frac{1}{\overline{w}_m}\frac{\partial}{\partial \hat{y}_m} w_m(x_f, y_f, \hat{x}_m, \hat{y}_m)|_{\hat{x}_m = x_m, \hat{y}_m = y_m}. \quad (1d)$$

where $\overline{w}_f = \overline{w}_m$ indicates the mean fitness of resident population with resource allocation strategy $x_f$, $y_f$, $x_m$ or $y_m$. Assuming no mutational correlation among the coevolving traits, the evolutionary dynamics of each evolving variable can be then captured by the canonical equation[7]:

$$\frac{dx_f}{dt} = \kappa \sigma_{x_f}^2 \beta_{x_f} \quad (2a)$$

$$\frac{dy_f}{dt} = \kappa \sigma_{y_f}^2 \beta_{y_f} \quad (2b)$$

$$\frac{dx_m}{dt} = \kappa \sigma_{x_m}^2 \beta_{x_m} \quad (2c)$$

$$\frac{dy_m}{dt} = \kappa \sigma_{y_m}^2 \beta_{y_m}. \quad (2d)$$

Here, the factor $\kappa = \frac{1}{2}\mu N$ represents the rate at which beneficial mutations contributing to evolutionary change arise per unit time. Specifically, $\mu$ is the mutation rate of the trait at birth, $N$ is the size of the resident population, and $\frac{1}{2}$ reflects that, under directional selection, only half of mutations are expected to be beneficial. The terms $\sigma_{x_f}^2$, $\sigma_{y_f}^2$, $\sigma_{x_m}^2$ and $\sigma_{y_m}^2$ denote the variances in the phenotypic effects of mutations for the four evolving traits.

Following Long and Henshaw's framework (equations 6 and 7 in ref. [6]), feedback effects on each selection gradient ($\beta_{x_f}$, $\beta_{y_f}$ $\beta_{x_m}$, $\beta_{y_m}$), arising from changes in resource allocation variables can be derived using the chain rule:

$$\frac{d}{dt}\beta_{x_f} = \frac{\partial \beta_{x_f}}{\partial x_f}\frac{dx_f}{dt} + \frac{\partial \beta_{x_f}}{\partial y_f}\frac{dy_f}{dt} + \frac{\partial \beta_{x_f}}{\partial x_m}\frac{dx_m}{dt} + \frac{\partial \beta_{x_f}}{\partial y_m}\frac{dy_m}{dt} \quad (3a)$$

$$\frac{d}{dt}\beta_{y_f} = \frac{\partial \beta_{y_f}}{\partial x_f}\frac{dx_f}{dt} + \frac{\partial \beta_{y_f}}{\partial y_f}\frac{dy_f}{dt} + \frac{\partial \beta_{y_f}}{\partial x_m}\frac{dx_m}{dt} + \frac{\partial \beta_{y_f}}{\partial y_m}\frac{dy_m}{dt} \quad (3b)$$

$$\frac{d}{dt}\beta_{x_m} = \frac{\partial \beta_{x_m}}{\partial x_f}\frac{dx_f}{dt} + \frac{\partial \beta_{x_m}}{\partial y_f}\frac{dy_f}{dt} + \frac{\partial \beta_{x_m}}{\partial x_m}\frac{dx_m}{dt} + \frac{\partial \beta_{x_m}}{\partial y_m}\frac{dy_m}{dt} \quad (3c)$$

$$\frac{d}{dt}\beta_{y_m} = \frac{\partial \beta_{y_m}}{\partial x_f}\frac{dx_f}{dt} + \frac{\partial \beta_{y_m}}{\partial y_f}\frac{dy_f}{dt} + \frac{\partial \beta_{y_m}}{\partial x_m}\frac{dx_m}{dt} + \frac{\partial \beta_{y_m}}{\partial y_m}\frac{dy_m}{dt}. \quad (3d)$$

We can then substitute equation $(2)$ into equation $(3)$. The resulting equations describe how changes in the resource allocation traits ($x_f$, $y_f$, $x_m$, $y_m$) affect the corresponding selection gradients ($\beta_{x_f}$, $\beta_{y_f}$ $\beta_{x_m}$, $\beta_{y_m}$), respectively, in the absence of mutational covariance. To illustrate how evolutionary changes in each trait feed back to modify the selection gradient, we take equation $(3a)$ as an example and focus on the selection gradient acting on the female trait $x_f$, which determines the relative allocation of resources to egg production and, consequently, the realised proportion of resource allocated to egg production $g_f$. Because four traits are coevolving, evolutionary change in each trait can potentially alter the selection environment experienced by $x_f$. Thus, the total feedback on can be decomposed into four components: (1) Direct selection on the focal trait $x_f$ can affect selection on the same trait ("within-trait feedback," $\frac{\partial \beta_{x_f}}{\partial x_f}\frac{dx_f}{dt} = \kappa\sigma_{x_f}^2\beta_{x_f}\frac{\partial \beta_{x_f}}{\partial x_f}$: green lines in Supplementary Figs. 12,13). (2) Direct selection on the coevolving trait expressed in the same sex, $y_f$, can affect the selection environment experienced by $x_f$ ("cross-trait feedback due to selection on the coevolving trait expressed in the same sex", $\frac{\partial \beta_{x_f}}{\partial y_f}\frac{dy_f}{dt} = \kappa\sigma_{y_f}^2\beta_{y_f}\frac{\partial \beta_{x_f}}{\partial y_f}$: purple lines in Supplementary Figs. 12,13). In addition, two other feedback components arise from direct selection on coevolving traits expressed in the opposite sex, here $x_m$ and $y_m$. Evolutionary change in these traits can alter the selection environment experienced by the focal trait $x_f$, giving rise to: (3) "cross-trait feedback due to selection on the gamete production strategy in the opposite sex" ($\frac{\partial \beta_{x_f}}{\partial x_m}\frac{dx_m}{dt} = \kappa\sigma_{x_m}^2\beta_{x_{m_f}}\frac{\partial \beta_{x_f}}{\partial x_m}$: yellow lines in Supplementary Figs. 12,13); and (4) "cross-trait feedback due to selection on the parental care strategy in the opposite sex" ($\frac{\partial \beta_{x_f}}{\partial y_m}\frac{dy_m}{dt} = \kappa\sigma_{y_m}^2\beta_{y_{m_f}}\frac{\partial \beta_{x_f}}{\partial y_m}$: orange lines in Supplementary Figs. 12,13).

To examine qualitatively how feedback affects selection, we consider the combination of the selection gradient and its feedback components[6]. A positive selection gradient indicates selection for trait values larger than the current resident strategy. If feedback is also positive, it strengthens this selection, making selection for larger trait values even stronger ("positive reinforcing" feedback). In contrast, if the selection gradient is positive but feedback is negative, feedback weakens selection for larger trait values ("positive inhibiting" feedback). Analogous considerations apply when the selection gradient is negative. Negative feedback on a negative selection gradient reinforces selection towards smaller trait values ("negative reinforcing" feedback), whereas positive feedback weakens selection towards smaller trait values ("negative inhibiting" feedback). Thus, feedback can either amplify or dampen selection on a trait, depending on whether it acts in the same direction as, or in the opposite direction to, the selection gradient. The overall feedback acting on a selection gradient is given by the sum of all feedback components. We can therefore compare the magnitude of each component to identify which pathway has the strongest effect on selection acting on the focal trait.

Supplementary Fig. 11 shows an example evolutionary trajectory derived from equation (2), in which males and females initially differ only in gamete size (with egg size $m_f = 0.1$ and sperm size $m_m = 0.01$), and each local group contains five males and five females. Starting from equal proportional allocation to parental care and gamete production in both sexes (with half of resources allocated to each reproductive activity), females evolve to allocate more resources to gamete production than males, whereas males evolve to allocate more resources to post-zygotic care than females. This trajectory converges to the same equilibrium as in Supplementary Fig. 10 when the same parameter settings are used.

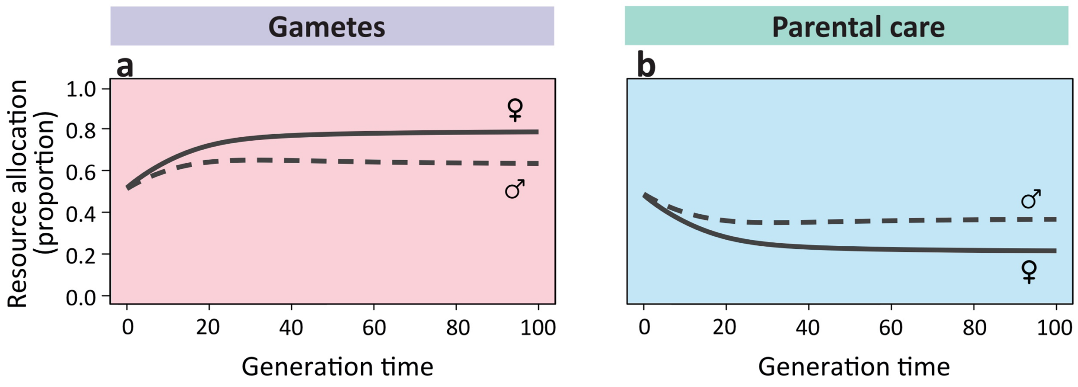


**Supplementary Figure 11. Evolutionary trajectories of resource allocation without mating competition.** The graphs depict representative time trajectories of resource allocation to **(a)** gamete production and **(b)** post-zygotic parental care, in the absence of any forms of pre-mating competition, with full certainty of paternity and non-depreciable parental care. When males and females initially differ only in gamete size ($m_f = 0.1, m_m = 0.01$), and both sexes start from equal proportional allocation to gamete production and parental care ($x_f = y_f = x_m = y_m = 0.1$, and thus $p_f = g_f = p_m = g_m = 0.5$), both sexes evolve increased allocation to gamete production. This increase is stronger in females, which ultimately allocate more resources to egg production than males allocate to sperm production. Conversely, both sexes decrease allocation to post-zygotic care, with females decreasing more, so that males ultimately provide more post-zygotic parental care. Key parameter values: $R = 1, k = 1, d = 10, r = 0.5$ (see Supplementary Table 1 for an overview of the notation).

Over evolutionary time, each selection gradient is shaped by within- and cross-trait feedback components (Supplementary Figs. 12, 13). Because female gametes are substantially larger than male gametes, the initial same resource investment produces far fewer eggs than sperm. Given the high fertilisation rate, the fitness value of eggs is approximately proportional to the number produced. Selection therefore favours females that allocate more resources to egg production (Supplementary Fig. 12$a_1$), shifting resources away from post-zygotic parental care (Supplementary Fig. 12$b_1$). Accordingly, the selection gradient for the egg production trait ($\beta_{x_f}$) remains positive (Supplementary Fig. 12$a_2$), whereas that for the female care trait ($\beta_{y_f}$) remains negative (Supplementary Fig. 12$b_2$) throughout the evolutionary trajectory, with both approaching zero as equilibrium is reached. Throughout this process, the four feedback components acting on $\beta_{x_f}$ are negative (Supplementary Fig. 12$a_3$ - $a_6$), corresponding to "positive inhibiting" feedback that weakens selection for increased resource allocation to egg production. Specifically, comparison of the absolute values of the four feedback components shows that within-trait feedback dominates during the initial phase of the evolution of female resource allocation to egg production (green background in Supplementary Fig. 12$a_3$). This arises because the initial allocation to egg production is already relatively high and the fitness function for egg production is concave. Consequently, further increases in allocation to egg production reduce the magnitude of the selection gradient favouring additional investment in egg production. Later in the trajectory, cross-trait feedback generated by selection on female parental care becomes dominant (purple background in Supplementary Fig. 12$a_4$). As females reduce their investment in parental care, offspring survival declines, which in turn weakens selection favouring further increases in egg production. This inhibiting cross-trait feedback remains dominant until the resource allocation strategy approaches equilibrium. The concurrent increase in allocation to egg production initially generates the dominant inhibiting feedback on selection for reduced female post-zygotic care (purple background in Supplementary Fig. 12$b_4$). As females allocate more resources to egg production, sufficient parental care is needed to ensure the survival of the additional offspring produced, thereby constraining further reductions in female care. Along with the progressive reduction in female care, however, within-trait feedback becomes dominant as the system approaches equilibrium (green background in Supplementary Fig. 12$b_3$). Overall, initial sex differences in gamete size cause females to invest more in egg production than in post-zygotic care through the trade-off between clutch size (i.e. egg number) and offspring survival.

The overall pattern of selection on male resource allocation strategies is partly analogous to the feedback dynamics described above for females, but important differences emerge (Supplementary Fig. 13). Because the local mating group size is relatively large (see Supplementary Fig. 10b), males experience strong sperm competition, favouring increased resource allocation to sperm production

(Supplementary Fig. 13$a_1$) and reduced allocation to post-zygotic care (Supplementary Fig. 13$b_1$). Accordingly, the selection gradient for allocation to sperm production is initially positive (Supplementary Fig. 13$a_2$), whereas that for male post-zygotic care is initially negative (Supplementary Fig. 13$b_2$). Similar to the pattern observed in females, self-inhibiting feedback initially dominates selection on sperm production (green background in Supplementary Fig. 13$a_3$). This is followed by a dominant role of cross-trait feedback generated by selection on male parental care (purple background in Supplementary Fig. 13$a_4$). Moreover, the increase in sperm allocation initially provides the strongest inhibiting feedback on selection for reduced post-zygotic care (purple background in Supplementary Fig. 13$b_4$). However, this does not fully explain the evolutionary dynamics of male resource allocation in this case, because changes in female allocation strategy also play a crucial role in shaping the selection environment experienced by male traits at later stages. In particular, as female care decreases, it becomes the dominant feedback component inhibiting both further increases in male sperm production and further decreases in male post-zygotic care (indicated by the orange background to the left of the grey dashed vertical lines in Supplementary Figs. 13$a_6$ and 13$b_6$, respectively). This dominant role persists until both selection gradients change sign, after which males begin to decrease allocation to sperm production and increase allocation to post-zygotic care (right of the grey dashed vertical line in Supplementary Fig. 13). After this switching point, declining female care reinforces selection for reduced sperm production and increased post-zygotic care, while remaining the dominant feedback component acting on both male traits until equilibrium is reached. Taken together, initial sex differences in gamete size cause males to invest more in sperm production than in post-zygotic care through the trade-off between sperm competition and offspring survival. However, the anisogamy-driven decrease in female care strongly inhibits both further increases in male sperm production and further decreases in male care. As a result, males increase allocation to gamete production less, and reduce allocation to parental care less, than females. This ultimately leads to an evolutionary outcome in which females allocate more resources to gamete production than males, whereas males provide more post-zygotic care than females.

In summary, these results provide a comprehensive understanding of the evolutionary process linking anisogamy to the subsequent distribution of post-zygotic care between the sexes. In the initial phase, both males and females allocate half of their resources to gamete production and half to post-zygotic care. Because females produce fewer gametes than males as a result of their larger gamete size, selection favours increased allocation to egg production in females, owing to the high fitness value of egg production when the fertilisation rate is high. Males also experience selection for increased allocation to gamete production due to sperm competition. In both sexes, these increases are initially shaped most strongly by within-trait feedback, which inhibits selection for further increases in gamete production because fitness returns are concave. This is followed by dominant cross-trait feedback arising from selection for reduced post-zygotic care within each sex, reflecting the allocation trade-off between offspring number and offspring survival.

In response to increased allocation to gamete production, both sexes reduce allocation to post-zygotic care. In particular, females reduce care more rapidly because selection for increased egg production is stronger, drawing more resources away from post-zygotic care compared with males. This reduction in care is initially strongly inhibited by increased allocation to gamete production in both sexes, because insufficient care would lead to low offspring survival. As female care declines more rapidly, it eventually reaches a level at which it becomes the dominant feedback component preventing further increases in male sperm production and further decreases in male care; otherwise, zygotes are unlikely to survive. This continues until selection on male traits changes direction, favouring reduced allocation to sperm production and increased allocation to post-zygotic care. After this switching point, declining female care continues to act as the dominant cross-trait feedback, reinforcing selection for reduced sperm production and increased male care until the system approaches equilibrium. As a result, females ultimately allocate more resources to gamete production, whereas males allocate more resources to post-zygotic care.

Our feedback analysis suggests a different perspective on Trivers' argument. In our model, the sex producing smaller gametes ultimately evolves to provide more post-zygotic care. This occurs because females produce relatively few, large gametes, so each additional egg has high fitness value when fertilisation efficiency is high. Selection therefore favours increased allocation to egg production, but this leaves fewer resources available for post-zygotic care. The resulting decline in female care then changes the selection environment experienced by males. Although males are initially selected to allocate more resources to sperm production because of sperm competition, reduced female care lowers offspring survival and thereby constrains both further increases in sperm production and further decreases in male care. Eventually, selection on males shifts towards reduced sperm production and increased post-zygotic care. Thus, sex differences in post-zygotic care emerge not from a self-reinforcing process in which 'one sex invests more because it has already invested more', but from via dynamic interactions among traits within and between the sexes.

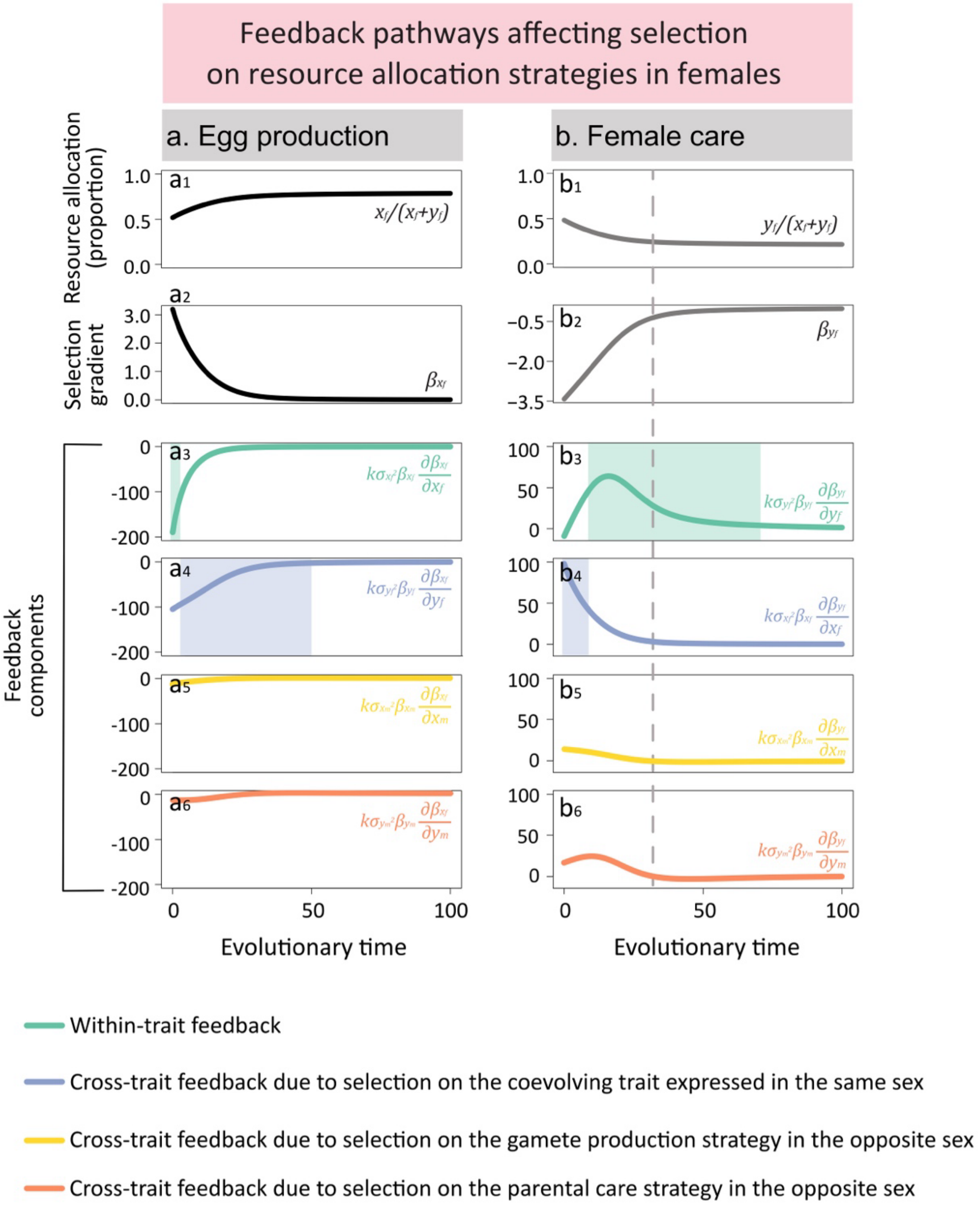


**Supplementary Figure 12. Feedback pathways linking anisogamy to selection on female traits**. As shown in Supplementary Fig. 11, anisogamy drives males to provide most post-zygotic care, while females invest more in egg production. Here, we use feedback analysis to examine how evolutionary changes in each of the four coevolving traits ($x_f, y_f, x_m, y_m$) feed back to shape selection on the others when the sexes initially differ only in gamete size. **($a_1$, $b_1$)** Evolutionary trajectories of $x_f$ and $y_f$, following the canonical equation of adaptive dynamics. **($a_2$, $b_2$)** Selection gradients $\beta_{x_f}$ and $\beta_{y_f}$ over evolutionary time. **($a_3$, $b_3$) "**Within-trait feedback" (green lines) on each selection gradient. **($a_4$, $b_4$)** "Cross-trait feedback due to selection on the coevolving trait expressed in the same sex" on each selection gradient (purple lines). **($a_5$, $b_5$)** "Cross-trait feedback due to selection on the gamete production strategy in the opposite sex" on each selection gradient (yellow lines). **($a_6$, $b_6$)** "Cross-trait feedback due to selection on the parental care strategy in the opposite sex" on each selection gradient (orange lines). The grey dashed vertical lines indicate the time points at which the male care trait transitions from a decreasing to an increasing trend. The corresponding background colours indicate which feedback component has the largest absolute value and therefore dominates among all feedback components. The population was initialised at $x_f = y_f = x_m = y_m = 0.1$, and thus $p_f = g_f = p_m = g_m = 0.5$; = 1 Other parameter values: $R = 1$, $m_f = 0.1$, $m_m = 0.01$, $k = 1$, $d = 10$, $r = 0.5$, $\theta_f = \theta_m = 1$, $\kappa = 1$ and $\sigma^2_{x_f} = \sigma^2_{y_f} = \sigma^2_{x_m} = \sigma^2_{y_m} = 1$. Evolutionary trajectories, selection gradients, and feedback components were approximated numerically using Wolfram Mathematica (see the Mathematica code in the Supplementary Materials for more detail).

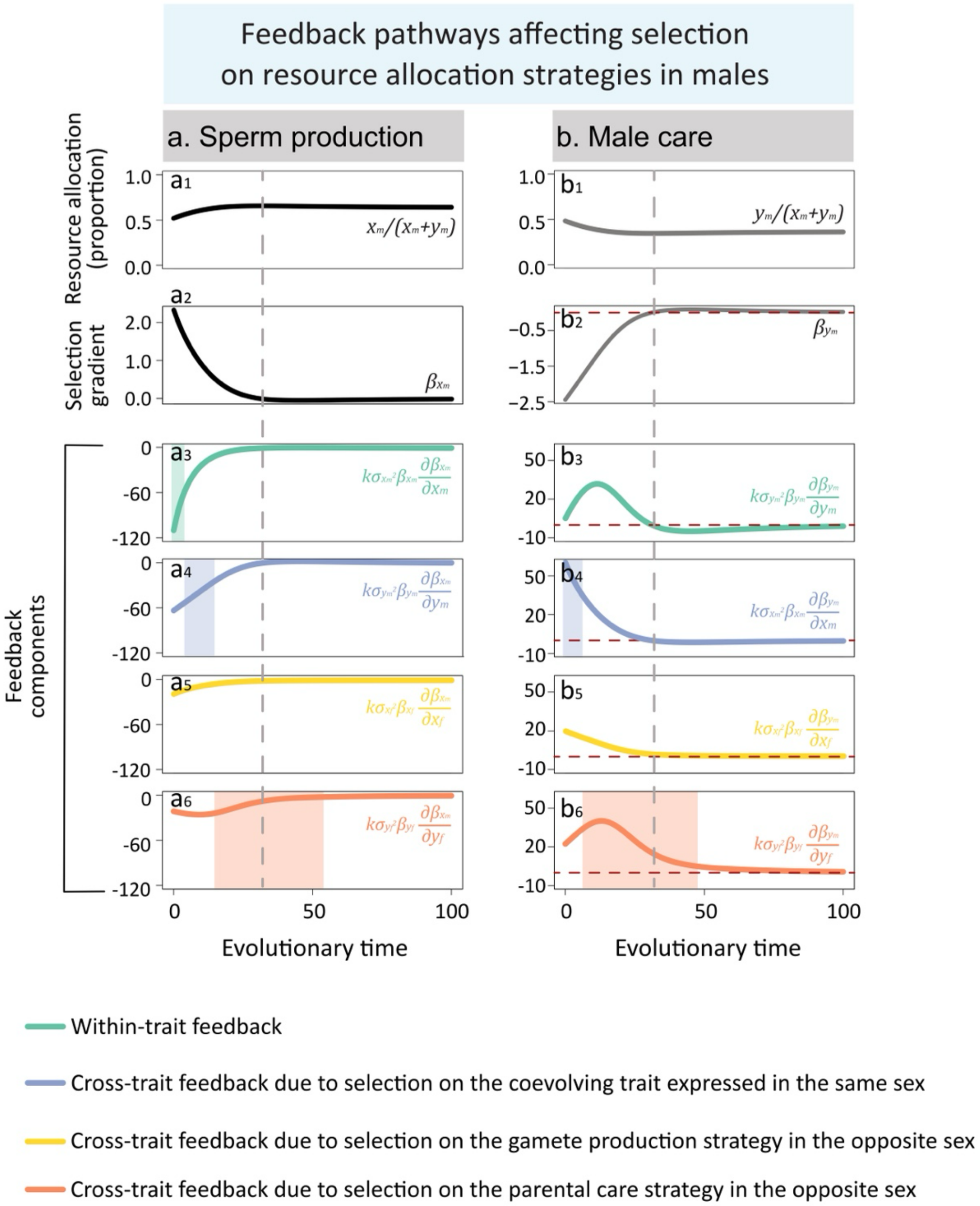


**Supplementary Figure 13. Feedback pathways linking anisogamy to selection on male traits.** This figure is analogous to Supplementary Fig. 12, but focuses on selection gradients and feedback components acting on male resource allocation trait, $x_m$ and $y_m$ . Specifically, it shows **($a_1$, $b_1$)** the evolutionary trajectories of male allocation to $x_m$ and $y_m$; **($a_2$, $b_2$)** the corresponding selection gradients $\beta_{x_m}$ and $\beta_{y_m}$; and **($a_3$-$a_6$, $b_3$-$b_6$)** the four feedback components acting on each selection gradient. Background colours indicate the feedback component with the largest absolute value and therefore the dominant feedback pathway. The grey dashed vertical lines indicate the time points at which the male care trait transitions from decreasing to increasing. The red dashed horizontal lines indicate the value of zero. Parameter values and numerical methods are as in Supplementary Fig. 12.

**Table 1. Overview of notation in the main text.** Subscripts $f$ and $m$ denote female and male, respectively. Traits of a mutant individual are indicated with a hat (^)

| Symbol | Meaning | Notes |
|---|---|---|
| **Resource allocation** | | |
| $R$ | Resource budget for each individual | Fixed parameter with a default value of 1 |
| $x_f, y_f, x_m, y_m$ | Evolving resource allocation strategies | Evolving parameters |
| $c_f, c_m$ | The fraction of resources allocated to mating competition | $c_f = x_f, c_m = x_m$ |
| $p_f, p_m$ | The fraction of resources allocated to parental care | $p_f = y_f(1 - x_f), p_m = y_m(1 - x_m)$ |
| $g_f, g_m$ | The fraction of resources allocated to gamete production | $g_f = 1 - c_f - p_f, g_m = 1 - c_m - p_m$ |
| **Gamete production** | | |
| $m_f, m_m$ | Gamete size (for eggs, sperm) | Fixed parameters with default values: $m_f = 0.1,\ m_m = 0.01$ |
| $n_{\text{egg}}$ | Number of eggs produced by a resident female | $n_{\text{egg}} = Rg_f/m_f$ |
| $n_{\text{sperm}}$ | Number of sperm produced by a resident male | $n_{\text{sperm}} = Rg_m/m_m$ |
| **Mating competition** | | |
| $k$ | Mate encounter efficiency (scramble competition scenario) | Fixed parameter with a default value of 1 |
| $d$ | Group size coefficient, representing the expected size of local groups when each individual is present in only one group (scramble competition scenario) | Fixed parameter with a default value of 10 |
| $r$ | Adult sex ratio | Fixed parameter with a default value of 0.5 |
| $\delta$ | "Winner-takes-all" effect (contest competition scenario) | Fixed parameter with a default value of 1 |
| $\theta_f, \theta_m$ | Average number of local groups a resident individual can enter | Under scramble competition, $\theta_f = kc_fR$ and $\theta_m = kc_mR$; under contest competition, $\theta_f$ and $\theta_m$ are fixed at 1 by default |
| $\theta'_f, \theta'_m$ | Expected number of local groups entered by resident females and males, respectively (≥1) | $\theta'_f = \frac{\theta_f}{1-e^{-\theta_f}}, \theta'_m = \frac{\theta_m}{1-e^{-\theta_m}}$ |
| $n_f, n_m$ | Number of resident females and males within each local group, respectively | $n_f = (1 - r)\theta_f d, n_m = r\theta_m d$ |
| $n'_{\text{egg}}$ | Number of eggs contributed to each local group by a resident female | $n'_{\text{egg}} = n_{\text{egg}}/\theta'_f$ |
| $n'_{\text{sperm}}$ | Number of eggs contributed to each local group by a resident male | $n'_{\text{sperm}} = n_{\text{sperm}}/\theta'_m$ |

| | | |
|---|---|---|
| $N_e, N_s$ | The total numbers of eggs and sperm within each local group, respectively | $N_e = n_f n'_{\text{egg}}, N_s = n_m n'_{\text{sperm}}$ |
| **Fertilisation within each local mating group** | | |
| $\gamma$ | Fertilisation efficiency | Fixed parameter with a default value of 1 |
| $F(N_e, N_s)$ | Total fertilisation within a local mating group | $F(N_e, N_s) = \frac{1+\gamma(N_e+N_s)-\sqrt{1+\gamma^2(N_e-N_s)^2+2\gamma(N_e+N_s)}}{2\gamma}$ |
| $f_f$ | Total number of fertilised eggs per resident female across local groups | $f_f = \frac{\theta_f}{n_f} F(N_e, N_s)$ |
| $f_m$ | Total number of fertilised sperm per resident male across local groups | $f_m = \frac{\theta_m}{n_m} F(N_e, N_s)$ |
| **Offspring survival** | | |
| $b$ | Baseline offspring survival without any post-zygotic care | Fixed parameter with a default value of 0 |
| $\alpha$ | The effectiveness of parental care in enhancing offspring survival | Fixed parameter with a default value of 1 |
| $O_f, O_m$ | The realised total number of offspring produced by a resident | $O_f = \frac{\theta'_f}{n_f} F(N_e, N_s), O_m = \frac{\theta'_m}{n_m} F(N_e, N_s)$ |
| $P_f, P_m$ | Proportions of resources allocated to maternal and paternal care that are received per offspring, respectively | When care is non-depreciable, $P_f = p_f$ $P_m = p_m$; when care is depreciable, $P_f = p_f/O_f, P_m = p_m/O_m$ |
| $S(P_f, P_m)$ | Offspring survival | $S(P_f, P_m) = 1 - \mathrm{e}^{-\left(b+\alpha R(P_f+P_m)\right)}$ |
| **Fitness of a resident** | | |
| $w_f, w_m$ | Fitness of resident population | $w_f = f_f S(P_f, P_m), w_m = f_m S(P_f, P_m)$ |
| **Invasion fitness under scramble competition** | | |
| $\hat{n}_{\text{egg}}$ | Number of eggs produced by a mutant female | $\hat{n}_{\text{egg}} = R\hat{g}_f/m_f$ |
| $\hat{n}_{\text{sperm}}$ | Number of sperm produced by a mutant male | $\hat{n}_{\text{sperm}} = R\hat{g}_m/m_m$ |
| $\hat{\theta}_f, \hat{\theta}_m$ | Average number of local groups a mutant can enter | $\hat{\theta}_m = k\hat{c}_m R$ , $\hat{\theta}_f = k\hat{c}_f R$ |
| $\hat{\theta}'_f, \hat{\theta}'_m$ | Expected number of local groups entered by a mutant female and a mutant male, respectively (≥1) | $\hat{\theta}'_f = \frac{\hat{\theta}_f}{1-e^{-\hat{\theta}_f}}$ , $\hat{\theta}'_m = \frac{\hat{\theta}_m}{1-e^{-\hat{\theta}_m}}$ |
| $\hat{n}'_{\text{egg}}$ | Number of eggs contributed to each local group by a mutant female | $\hat{n}'_{\text{egg}} = \hat{n}_{\text{egg}}/\hat{\theta}'_f$ |
| $\hat{n}'_{\text{sperm}}$ | Number of sperm contributed to each local group by a mutant male | $\hat{n}'_{\text{sperm}} = \hat{n}_{\text{sperm}}/\hat{\theta}'_m$ |
| $\hat{N}_e$ | Total number of eggs from resident and a mutant female within a local group | $\hat{N}_e = \hat{n}'_{\text{egg}} + (n_f - 1)n'_{\text{egg}}$ |

| | | |
|---|---|---|
| $\hat{N}_s$ | Total number of sperm from resident and a mutant male within a local group | $\hat{N}_s = \hat{n}'_{\text{sperm}} + (n_m - 1)n'_{\text{sperm}}$ |
| $\hat{f}_f$ | Total number of fertilised eggs produced by a mutant female across local groups | $\hat{f}_f = \hat{\theta}_f \left(\frac{\hat{n}'_{\text{egg}}}{\hat{N}_e}\right) F(\hat{N}_e, N_s)$, |
| $\hat{f}_m$ | Total number of fertilised sperm produced by a mutant male across local groups | $\hat{f}_m = \hat{\theta}_m \left(\frac{\hat{n}'_{\text{sperm}}}{\hat{N}_s}\right) F(N_e, \hat{N}_s)$ |
| $\hat{w}_f, \hat{w}_m$ | Invasion fitness of a mutant female and a mutant male, respectively | $\hat{w}_f = \hat{f}_f S(\hat{P}_f, P_m)$, $\hat{w}_m = \hat{f}_m S(P_f, \hat{P}_m)$ |
| **Invasion fitness under contest competition** | | |
| $\hat{n}'_f$ , $\hat{n}'_m$ | Effective number of rivals for a mutant female and a mutant male, respectively | $\hat{n}'_f = (n_f - 1)\left(\frac{c_f}{\hat{c}_f}\right)^{\delta}$, $\hat{n}'_m = (n_m - 1)\left(\frac{c_m}{\hat{c}_m}\right)^{\delta}$ |
| $\hat{N}'_e$ | Total number of effective eggs within a local group | $\hat{N}'_e = \hat{n}'_{\text{egg}} + \hat{n}'_f n'_{\text{egg}}$, where $\hat{n}'_{\text{egg}}$ and $n'_{\text{egg}}$ denotes the number of eggs contributed to each local group by a mutant and resident female, respestively |
| $\hat{N}'_s$ | Total number of effective sperm within a local group | $\hat{N}'_s = \hat{n}'_{\text{sperm}} + \hat{n}'_m n'_{\text{sperm}}$, where $\hat{n}'_{\text{sperm}}$ and $n'_{\text{sperm}}$ denotes the number of sperm contributed to each local group by a mutant and resident male, respestively |
| $\hat{f}'_f$ | Total number of fertilised eggs produced by a mutant female across local groups | $\hat{f}'_f = \theta_f \left(\frac{\hat{n}'_{\text{egg}}}{\hat{N}'_e}\right) F(\hat{N}'_e, N_s)$, |
| $\hat{f}'_m$ | Total number of fertilised sperm produced by a mutant male across local groups | $\hat{f}'_m = \theta_m \left(\frac{\hat{n}'_{\text{sperm}}}{\hat{N}'_s}\right) F(N_e, \hat{N}'_s)$ |
| $\hat{w}_f, \hat{w}_m$ | Invasion fitness of a mutant | $\hat{w}_f = \hat{f}'_f S(\hat{P}_f, P_m)$, $\hat{w}_m = \hat{f}'_m S(P_f, \hat{P}_m)$ |
| **Invasion fitness in the presence of uncertain paternity** | | |
| $\eta$ | The degree of uncertain paternity | Ranging from 0 to 1 |
| $\hat{w}_m$ | Invasion fitness of a mutant male | $\hat{w}_m = (1-\eta)\hat{f}_m S(P_f, \hat{P}_m) + \eta \hat{f}_m S(P_f, P_m)$ under scramble competition; $\hat{w}_m = (1-\eta)\hat{f}'_m S(P_f, \hat{P}_m) + \eta \hat{f}'_m S(P_f, P_m)$ under contest competition |
| **Invasion fitness with depreciable parental care** | | |
| $\hat{O}_f$ | The realised total number of offspring produced by a mutant female | $\hat{O}_f = \hat{\theta}'_f \frac{\hat{n}'_{\text{egg}}}{\hat{N}'_e} F(\hat{N}'_e, N_s)$ under contest competition; $\hat{O}_f = \hat{\theta}'_f \frac{\hat{n}'_{\text{egg}}}{\hat{N}_e} F(\hat{N}_e, N_s)$ under scramble competition |

| | | |
|---|---|---|
| $\hat{O}_m$ | The realised total number of offspring produced by a mutant male | $\hat{O}_m = \hat{\theta}_m' \frac{\hat{n}_{\text{sperm}}'}{\hat{N}_s'} F(N_e, \hat{N}_s')$ under contest competition; $\hat{O}_m = \hat{\theta}_m' \frac{\hat{n}_{\text{sperm}}'}{\hat{N}_s} F(N_e, \hat{N}_s)$ under scramble competition |
| $O_f'$ | The realised total number of offspring produced by a resident female | $O_f' = \frac{\theta_f' - 1}{n_f} F(N_e, N_s) + \frac{1}{n_f} F(N_e, \hat{N}_s')$ under contest competition; $O_f' = \frac{\theta_f' - 1}{n_f} F(N_e, N_s) + \frac{1}{n_f} F(N_e, \hat{N}_s)$ under contest competition |
| $O_m'$ | The realised total number of offspring produced by a resident male | $O_m' = \frac{\theta_m' - 1}{n_m} F(N_e, N_s) + \frac{1}{n_m} F(\hat{N}_e', N_s)$ under contest competition; $O_m' = \frac{\theta_m' - 1}{n_m} F(N_e, N_s) + \frac{1}{n_m} F(\hat{N}_e, N_s)$ under contest competition |
| $\hat{P}_f, \hat{P}_m$ | Per-offspring parental care provided by a mutant parent | $\hat{P}_f = \hat{p}_f / \hat{O}_f, \hat{P}_m = \hat{p}_m / \hat{O}_m$ |
| $P_f, P_m$ | Per-offspring parental care provided by a resident parent | $P_f = p_f / O_f', P_m = p_m / O_m'$ |

### References


1. Henshaw, J. M., Marshall, D. J., Jennions, M. D. & Kokko, H. Local gamete competition explains sex allocation and fertilization strategies in the sea. *Am. Nat.* **184**, E32–E49 (2014).
2. Henshaw, J. M., Bittlingmaier, M. & Schärer, L. Hermaphroditic origins of anisogamy. *Philos. Trans. R. Soc. B Biol. Sci.* **378**, 20220283 (2023).
3. Lehtonen, J. & Kokko, H. Two roads to two sexes: unifying gamete competition and gamete limitation in a single model of anisogamy evolution. *Behav. Ecol. Sociobiol.* **65**, 445–459 (2011).
4. Lehtonen, J., Parker, G. A. & Schärer, L. Why anisogamy drives ancestral sex roles. *Evolution* **70**, 1129–1135 (2016).
5. Henshaw, J. M., Jones, A. G. & Schärer, L. Anisogamy explains why males benefit more from additional matings. *Nat. Commun.* **13**, 3893 (2022).
6. Long, X., Lehtonen, J. & Henshaw, J. M. Quantifying feedback among traits in coevolutionary models. *Evolution* **79**, 2597–2611 (2025).
7. Dieckmann, U. & Law, R. The dynamical theory of coevolution: a derivation from stochastic ecological processes. *J. Math. Biol.* **34**, 579–612 (1996).
